# Colorimetric detection methods of pH-sensing wound dressing for point-of-care wound diagnostics

**DOI:** 10.1101/2024.10.11.617850

**Authors:** Katia Cherifi, Farnoush Toupchinejad, Katerina Christodoulopoulos, Aylin Kizilkaya, Simon Matoori

**Affiliations:** Faculté de Pharmacie, Université de Montréal, Montreal, QC H3T 1J4, Canada; Department of Pharmacology and Therapeutics, McGill University, Montreal, QC H3A 0G4, Canada

**Keywords:** Colorimetry, diagnostics, smartphone, wound pH, chronic wounds, Arduino

## Abstract

Chronic lower extremity wounds like diabetic foot ulcers (DFUs) are a major complication of diabetes and the leading cause of lower limb amputations worldwide. Currently, DFUs are diagnosed through macroscopic evaluation, and molecular diagnostics are lacking for the disease staging, treatment selection, and evaluation of treatment success. There is a need for new diagnostic technologies combined with detection methods for point-of-care use. Preclinical and clinical data support the importance of wound pH as a biomarker in chronic wound healing, with DFUs typically exhibiting more alkaline pH values when compared to normal healing wounds. In a previous study, we developed a pH-sensing fluorescent bandage based on pyranine-loaded microparticles that were physically immobilized in an alginate hydrogel. Here, we present a second-generation pH-sensing bandage for colorimetric wound diagnostics. Pyranine was adsorbed at high concentrations onto microparticles to enable colorimetric signal detection. This approach leverages pyranine’s ability to change color in response to pH variations through proton exchange properties with the wound fluid. The colorimetric properties of our bandage enable signal detection by two methods suited for point-of-care use: a smartphone camera and a cost-effective homebuilt RGB detector. Analyzing the color intensity of the bandage with Red, Green and Blue absorbance values, it is possible to correlate the RGB absorbance to a pH value in the clinically-relevant range to 6.0 to 9.0 *in vitro* and *ex vivo*, as the B values decreased with the increase in pH levels, as associated with DFUs. These findings indicate the potential of colorimetric detection using smartphone cameras or home-built absorbance detectors for rapid wound diagnostics at the point-of-care.

**Graphical Abstract:** A pH-sensing bandage enables colorimetric detection of chronic wounds. The bandage absorbs wound exudate, triggering proton exchange with dye-loaded microparticles in an alginate matrix. Chronic wounds with high pH values induce strong fluorescence and yellow coloration. The signal is quantified through RGB colorimetry analysis using a smartphone camera or a homebuilt RGB detector, enabling point-of-care diagnostics for chronic wounds.

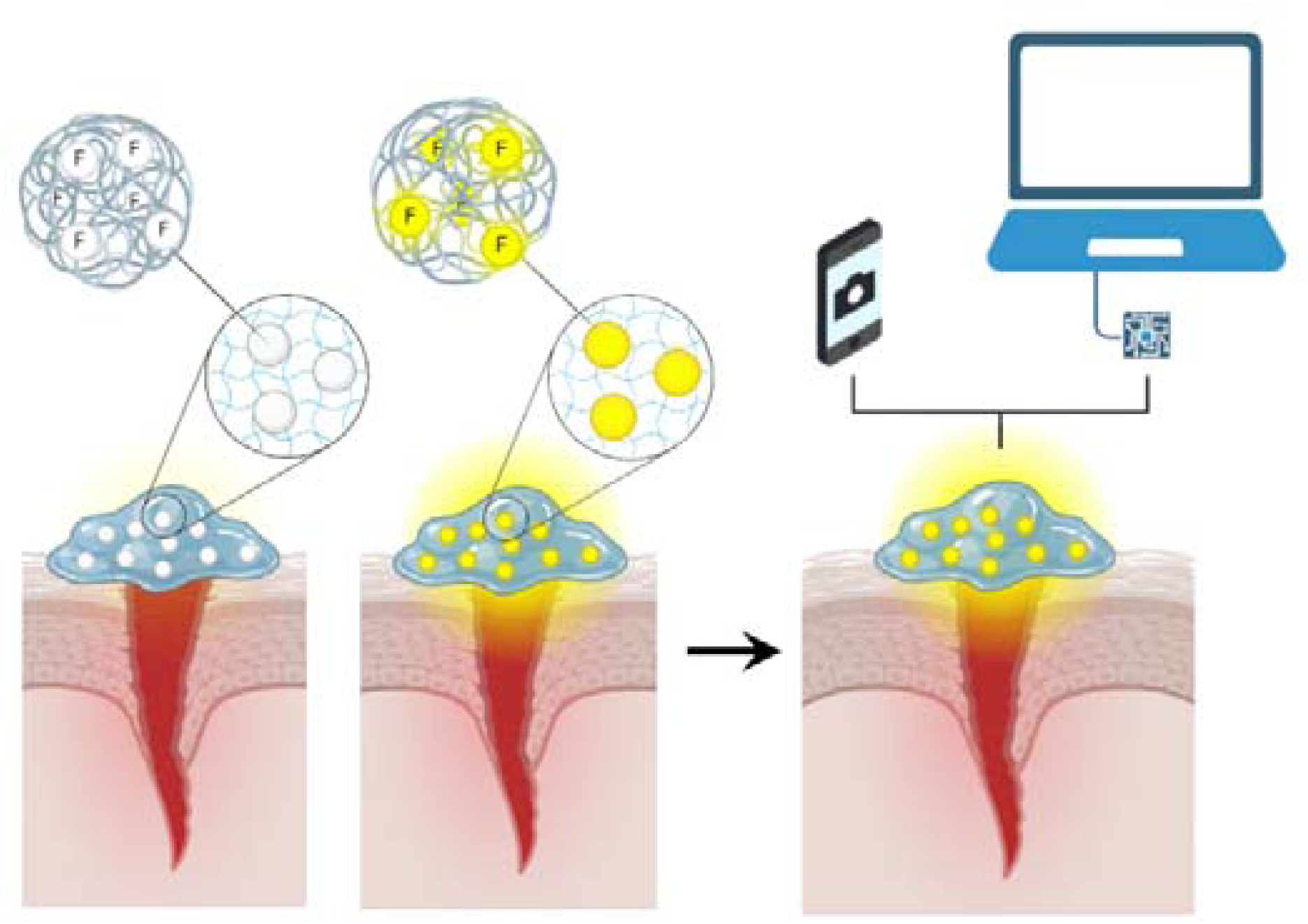

## 1. Introduction

Chronic lower extremity wounds (diabetic foot ulcers, *DFUs*) are a prevalent and serious complication of diabetes that affects up to 34% of diabetics ^[1]^. The healing rate is low as only half of these wounds fully heal within 7 months, and the recurrence rate is high with up to 60% after 3 years. DFUs are the leading cause of lower limb amputation worldwide, affecting 20% of patients^[2a-e]^. Despite our growing understanding of the molecular pathophysiology of diabetic wounds, current clinical guidelines recommend macroscopic evaluation of the wound based on ulcer depth, the implication of other tissues, and non-specific inflammatory markers such as redness and swelling ^[3a-c]^. Considering the lack of molecular diagnostics for DFU staging and prognosis, there is high interest in developing new point-of-care technologies for chronic wound diagnosis and monitoring. With the advent of molecular treatments for DFU, molecular diagnostics may also be useful to select the most suited treatment for individual patients with DFUs, and to evaluate treatment success.

Previous studies have highlighted the role of wound pH as a therapeutic target in chronic wounds ^[4a-d]^. It has been shown that chronic wounds such as diabetes tend to exhibit a more alkaline wound pH (7.5 to 9.0) when compared to normal healing wounds, which tend to have a slightly acidic pH approaching that of normal skin flora (around pH 6.0) ^[5a]^. The higher and more alkaline pH found in diabetic wounds reduces wound oxygenation, increases proteolytic enzyme activity, promotes the polarization of macrophages to the pro-inflammatory phenotype (M1), and promotes the growth of pathogenic bacteria ^[4a-d]^. An acidic to neutral pH reduces chronic inflammation and bacterial growth, and promotes macrophage polarization to the pro-healing phenotype (M2) ^[5b-e]^. A low wound pH creates favorable conditions for pro-healing processes such as angiogenesis and the migration and proliferation of keratinocytes and fibroblasts ^[6]^. Wound pH is also distributed heterogeneously in the wound bed, which makes it an interesting target for real-time and spatial monitoring of wound healing ^[7]^.

In a previous study ^[8]^, we developed a fluorescent pH-sensing bandage for point-of-care wound pH detection. This system is based on a double-encapsulation method to minimize dye leakage into the wound tissue: a pH-sensitive dye is loaded onto microparticles, and the microparticles are immobilized in a hydrogel matrix. The anionic pH-sensitive pyranine dye (HPTS) is loaded onto cationic microparticles, and the dye-loaded microparticles are encapsulated into ionically crosslinked calcium alginate hydrogels. As wound fluid enters the hydrogel, the fluorescence profile of the dye changes in relation to wound pH. We tested this diagnostic hydrogel on full excisional mouse wounds, and observed an increase in fluorescence with increasing pH. In a clinical setting, the fluorescence signal will be quantified with a portable fluorometer for point-of-care pH sensing. Considering the fact that portable fluorometers are emergent technologies with limited availability and comparably high costs, there is a need to develop diagnostic systems with alternative detection methods.

Potentiometric devices have been developed for pH sensing but have disadvantages because of their toxic components, the need for a battery, or their capacity to only assess pH at a single point. Colorimetric-based sensors based on pH indicators are simple and reliable systems that allow fast pH measurements due to their rapid proton exchange properties. In this study, we exploited the colorimetric properties of pyranine (the pH sensor of the fluorescent pH-sensing bandage) at higher concentrations to enable pH quantification through RGB colorimetry. The RGB color model is an additive color system used in digital displays and imaging. On electronics, colors are produced by adding light and by mixing different intensities of red, green and blue ^[9a-b]^. Pyranine’s absorbance properties change from colorless/slightly yellow at acidic pH to bright yellow at alkaline pH with a pKa of approx. 7.5 ^[8]^. Based on the pH-dependent absorbance profile of pyranine, we aimed to develop a colorimetric second generation of the pH-sensitive bandage and to test this system with two point-of-care detection methods: a smartphone camera and a cost-effective homebuilt RGB detector. The use of a smartphone camera as biodetection system allows for point-of-care measurements and is more widely available to the general public. The use of a homebuilt RGB detector offers a cost-effective and accessible solution for remote populations without compromising analytical performance; it can be easily assembled and used in resource-limited settings, ensuring equal access to healthcare.

### 2. Results & Discussion

### 2.1. ColorMeter-based detection enabled by smartphone camera

#### 2.1.1. Detection of HPTS using smartphone camera

To investigate if the absorbance of pyranine (HPTS) can be quantified by a smartphone camera, we prepared different dye concentrations and tested a commercial RGB-quantifying app (ColorMeter RGB Colorimeter) using a smartphone (iPhone 13). The pyranine solutions at different concentrations exhibited visibly different color intensities (**Supplementary Information Figure S1A**). As the dye concentration increased, the app detected a decrease in the blue signal intensity, reaching saturation at around 100 µM (**Figure 1B**). We used the blue channel as it detects signals between 400 and 500 nm, where pyranine exhibits a strong absorbance signal (**Figure 1A**). The decrease in blue channel signal intensity is due to the absorption of these photons by the dye. To investigate if the pH-dependence of HPTS can be detected by the app, we incubated it at pathophysiologically relevant pH values of 6.0 to 9.0 and tested the RGB-quantifying app. An increase in color intensity was visible as the pH value increased (**Supplementary Information Figure S1B**). This visible increase in color intensity was observed as a decrease in the blue signal intensity (**Figure 1C**).

**Figure 1.**
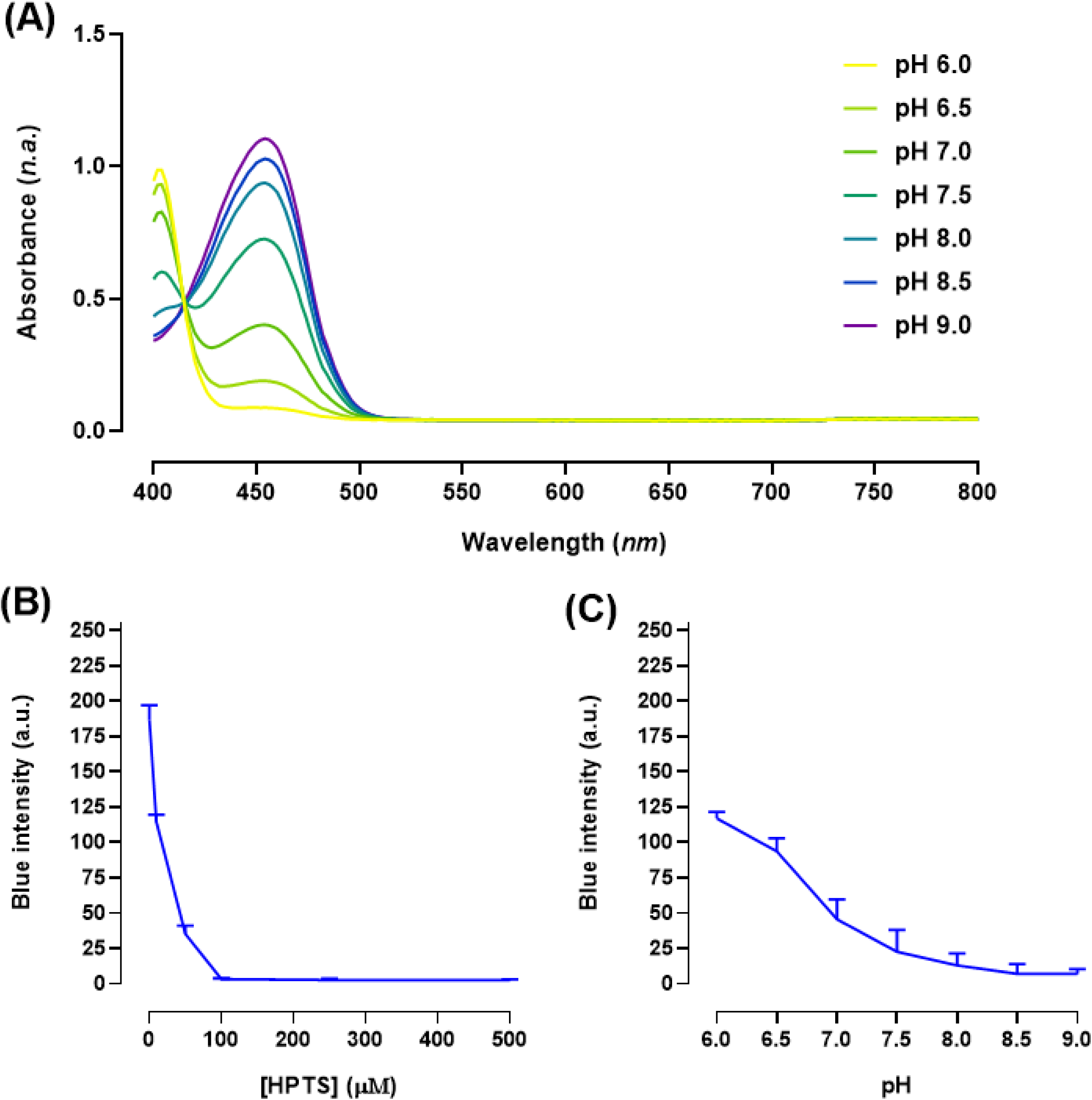
Investigation of the detection of pyranine (HPTS) in solution. **(A)** Absorbance spectrum of the dye. Pyranine was reconstituted in PBS isotonic buffer 50 mM at pH 6.0-9.0 and absorbance spectrum was measured between 400-800 nm with a Spark Tecan plate reader. **(B)** ColorMeter-recorded RGB absorbance of HPTS solutions at different concentrations in PBS isotonic buffer (50 mM, pH 7.4). **(C)** ColorMeter-recorded RGB absorbance of HPTS solutions at different pH values in MES isotonic buffer 50 mM for pH 6.0-6.5 and TBS isotonic buffer 50 mM for pH 7.0-9.0. Results presented as mean ± SD (n=3).

#### 2.1.2. Detection of HPTS-loaded BTA MPs using smartphone camera

To investigate if pyranine loaded onto microparticles can be detected by a smartphone camera, we incubated BTA microparticles with the dye and quantified the color intensity of this complex by using the same app as in *2.1.1*. As absorbance meters are less sensitive than fluorescence detectors, we used 13.3-times higher dye concentrations than with the fluorescence-based hydrogel ^[8]^. We observed a decrease in blue channel intensity when the MPs were loaded at increasing HPTS concentrations at pH 7.0 (**Figure 2A**). To investigate if these HPTS-loaded MPs can be detected by the app, we incubated them at pathophysiologically relevant pH values of 6.0 to 9.0. We observed visible color change as the pH increased (**Supplementary Information Figure S2**) and a decrease in blue channel intensity (**Figure 2B**). We selected a concentration of 100 µM for the following experiments because of the saturation of the signal at 500 µM.

**Figure 2.**
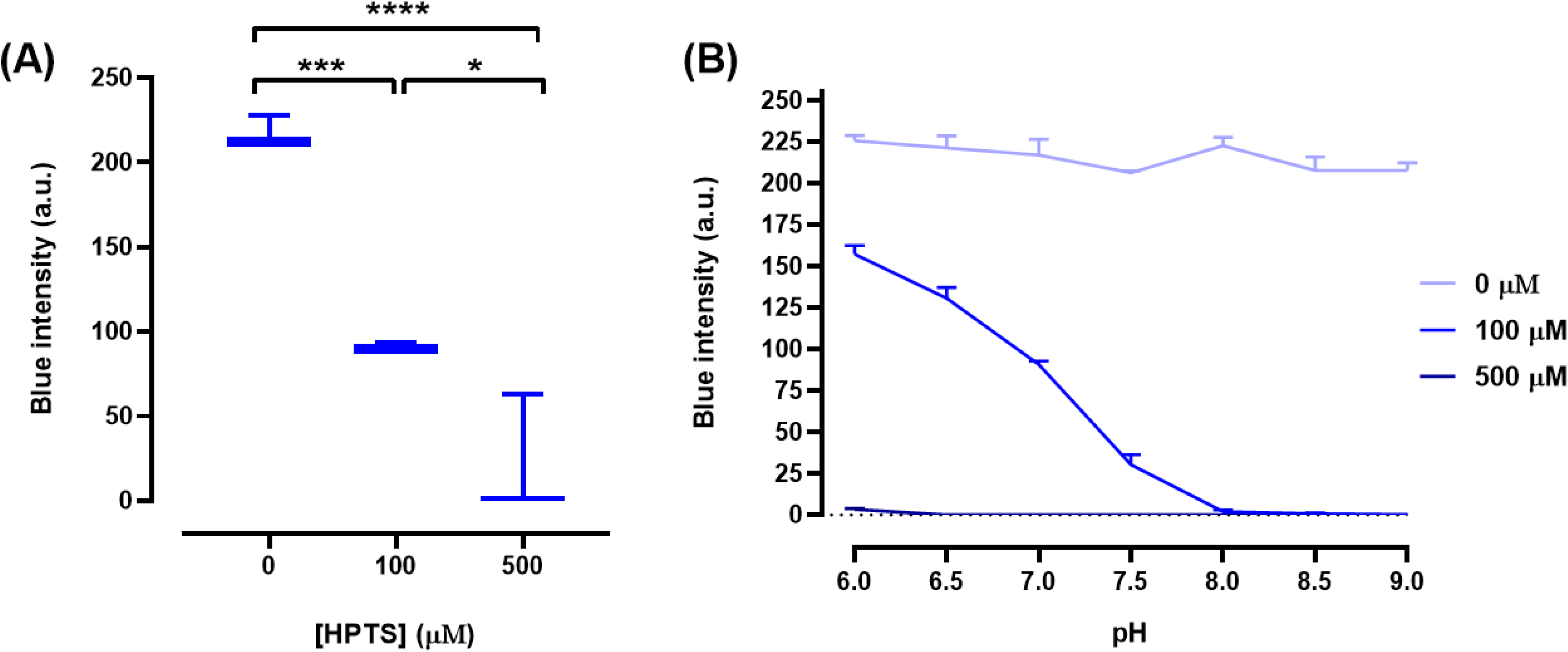
ColorMeter-recorded RGB absorbance of HPTS-loaded microparticles. **(A)** Concentration dependence of the signal in pyranine-loaded MPs at pH 7.0. **(B)** MPs loaded with different concentrations of HPTS at pH 6.0-9.0; Results presented as mean ± SD (n=3). Statistical significance was determined by one-way ANOVA followed by Tukey’s post-hoc test. *p <0.05, *** <0.001 **** <0.0001.

#### 2.1.3. Detection of pH-sensitive hydrogels using smartphone camera

To investigate if the pH-dependent color signal of the dye-loaded MPs can be detected in a hydrogel, we encapsulated them in a calcium-crosslinked alginate hydrogel and incubated them at pH 6.0 to 9.0 in corresponding buffer solutions. In order to detect colorimetric signal all over the bandage, we adapted the hydrogel formulation to increase the quantity of MPs of our initial formulation ^[8]^ from 5 mg to 150 mg (**Supplementary Information Figure S3**). We incubated the hydrogels in buffer solutions at pH 6.0 to 9.0 and quantified the color intensity as previously described with the smartphone app both *in vitro* in 6-well plates (**Supplementary Information Figure S4**) and *ex vivo* on full-thickness excisional dorsal mouse wounds (**Supplementary Information Figure S5**). We observed a decrease in blue intensity as pH values increased, both *in vitro* (**Figure 3A**) and *ex vivo* (**Figure 3B**); the similarity in the Blue channel decreases demonstrates that the measurement of RGB absorbance shows little interference from the background colors of an open wound, likely because of the high density of microparticles in the hydrogel.

**Figure 3.**
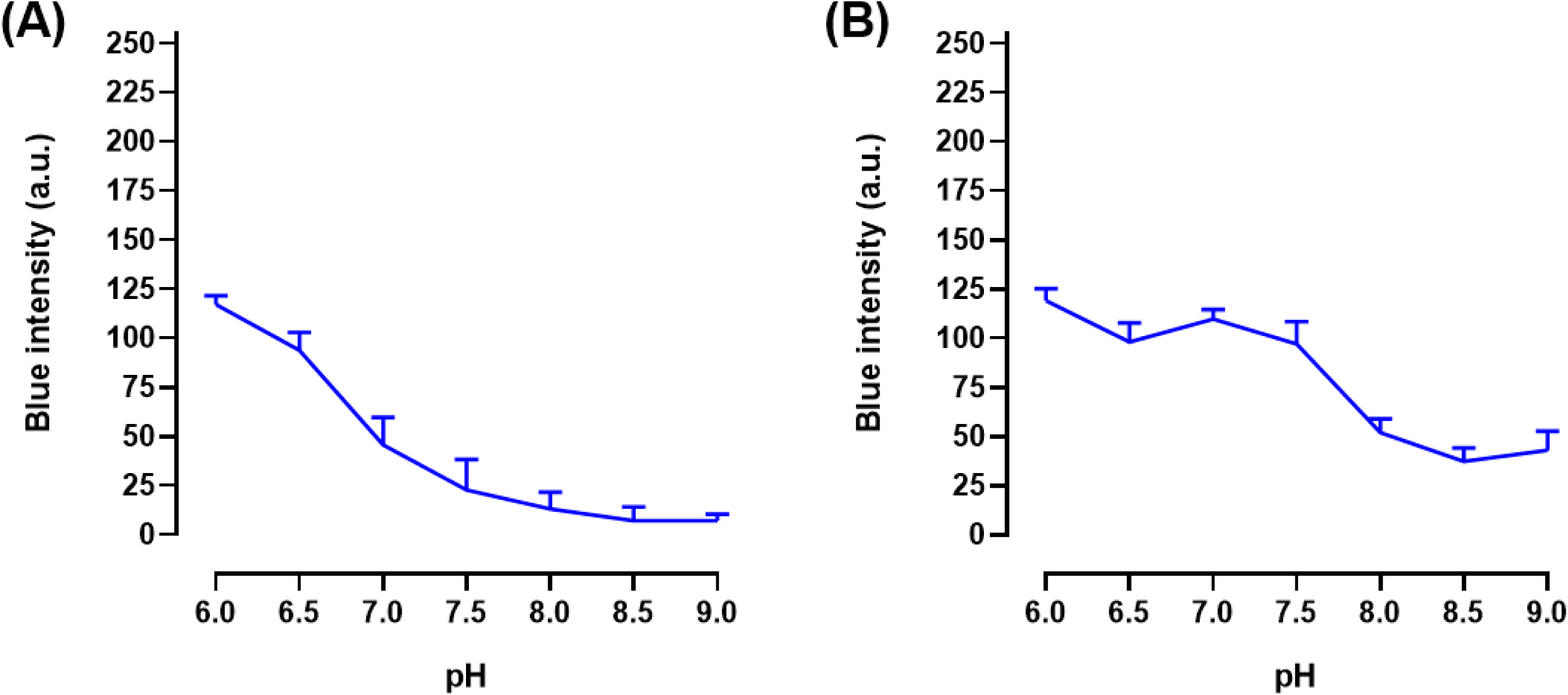
pH sensitivity of pyranine-loaded benzyltrialkylammonium (BTA) microparticles encapsulated in calcium alginate hydrogels. **(A)** ColorMeter-recorded blue signal intensity of hydrogels at different pH values in vitro. Hydrogels were incubated in MES isotonic buffer (50 mM) for pH 6.0-6.5 and TBS isotonic buffer (50 mM) for pH 7.0-9.0 for approx. 5 minutes before measurement. **(B)** Blue signal intensity of hydrogels at different pH values in full-thickness excisional mouse wounds *ex vivo*. Results presented as mean ± SD (n=3).

To determine sensing kinetics, we incubated the hydrogels in buffers of different pH values for up to 120 min. A visible distinction in signal between pH values was observed almost instantaneously; the color change at different pH values was visible at 2 min (**Figure S6**). A clear distinction in blue channel signal intensity can be seen at 3 min (**Figure 4**). As the protons diffused further in the pores of the hydrogel matrix, the yellow color of the bandages became more intense at higher pH values and blue channel signal intensity plateaued at approximately 30 min.

**Figure 4.**
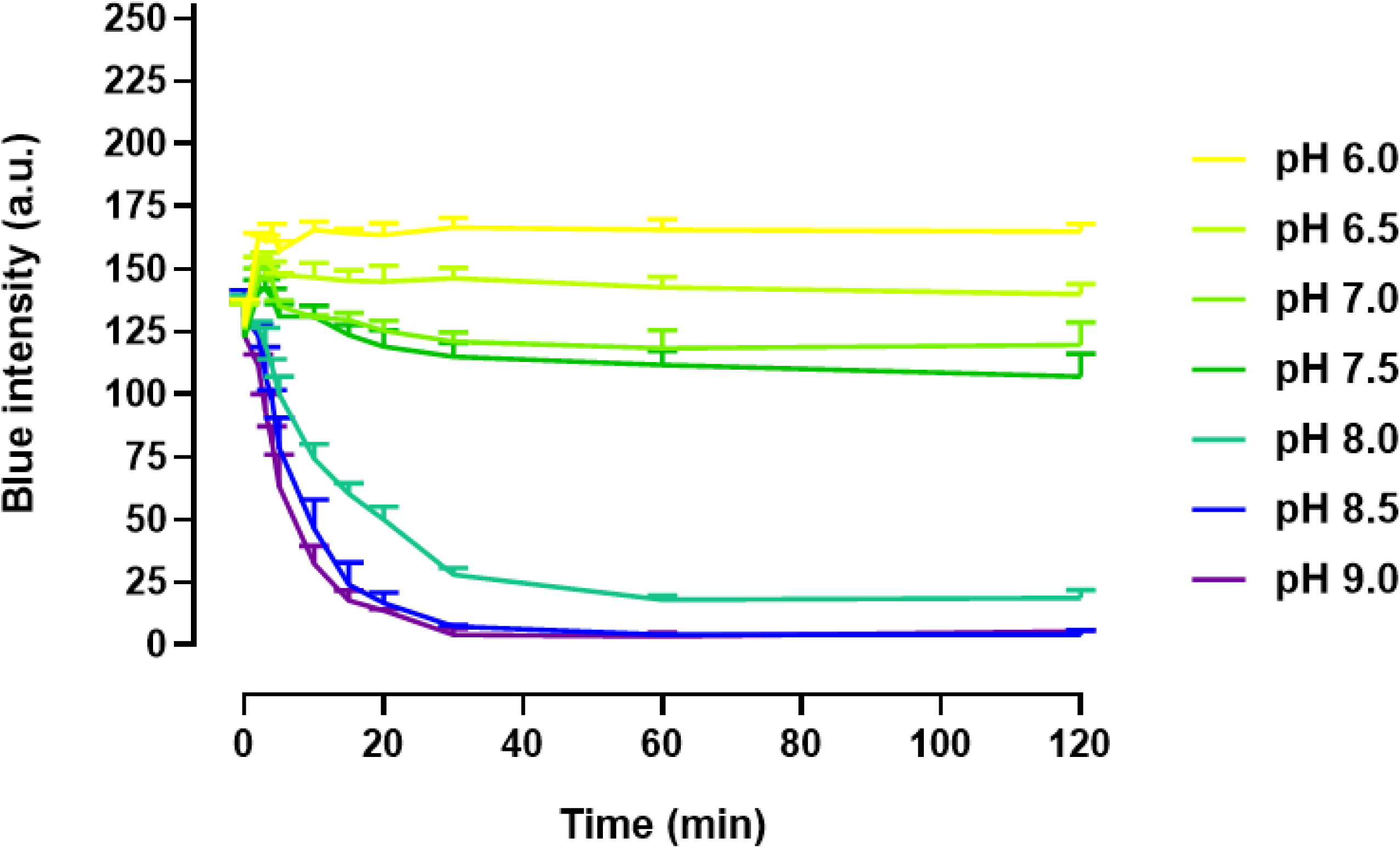
Kinetics of pH-sensing hydrogels *in vitro.* ColorMeter-recorded blue signal intensity of pyranine-loaded BTA microparticle-encapsulating calcium alginate hydrogels over time at different pH values. Hydrogels were incubated in MES isotonic buffer (50 mM) for pH 6.0-6.5 and TBS isotonic buffer (50 mM) for pH 7.0-9.0. Results presented as mean ± SD (n=3).

### 2.2. Arduino-coupled RGB sensor for colorimetry

#### 2.2.1. Detection of HPTS by Arduino-coupled RGB sensor

In the second part of this study, we developed a homemade absorbance sensor based on the Arduino technology coupled to a commercial RGB sensor. To investigate if the absorbance of pyranine solutions can be quantified by this detector, we prepared different dye concentrations. The solutions at different concentrations exhibited visibly different color intensities (Figure S1A). As the dye concentration increased, the sensor detected a decrease in the blue signal intensity, reaching saturation at around 500 µM (**Figure 5A**). To investigate if the pH-dependance of HPTS can be detected and quantified with this homemade RGB sensor, we incubated pyranine at pH values from 6.0 to 9.0 as described in *2.1.1.* and analyzed the signals with the Arduino-based sensor. The increase in pH value can be correlated with a decrease in the Blue channel signal (**Figure 5B**).

**Figure 5.**
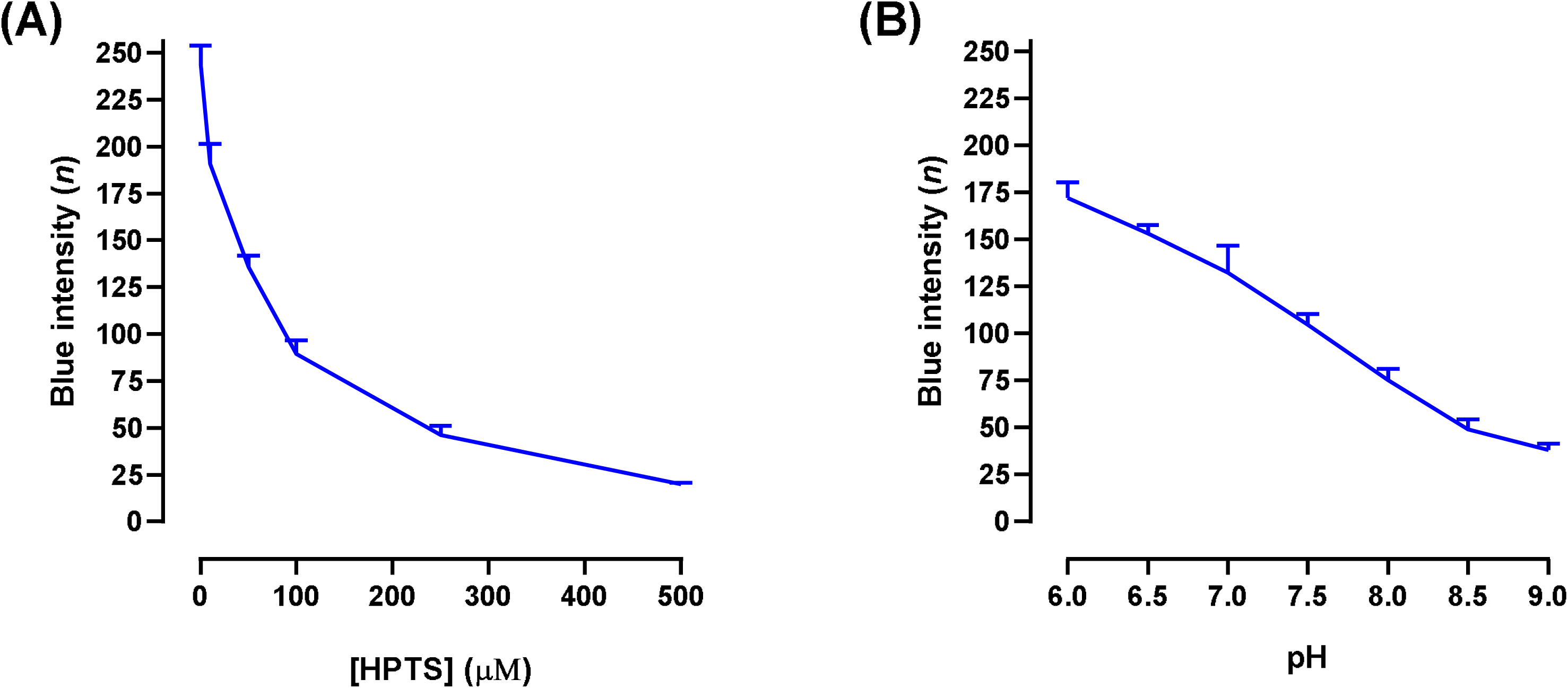
Arduino-recorded RGB absorbance of pyranine (HPTS) solutions. **(A)** RGB absorbance of HPTS at different concentrations in PBS isotonic buffer (50 mM, pH 7.4). **(B)** RGB absorbance of HPTS at different pH values in MES isotonic buffer (50 mM) for pH 6.0-6.5 and TBS isotonic buffer (50 mM) for pH 7.0-9.0. Results presented as mean ± SD (n=3).

#### 2.2.2. Detection of HPTS-loaded BTA MPs using Arduino-coupled RGB detector

To investigate if pyranine loaded onto microparticles can be detected by the Arduino-coupled RGB detector, we incubated BTA microparticles with pyranine and quantified the color intensity between pH values of 6.0 to 9.0. We observed a decrease of the B channel signal intensity in this pH range (**Figure 6**). The Arduino-coupled RGB sensor showed a similar decrease as the smartphone camera detection system.

**Figure 6.**
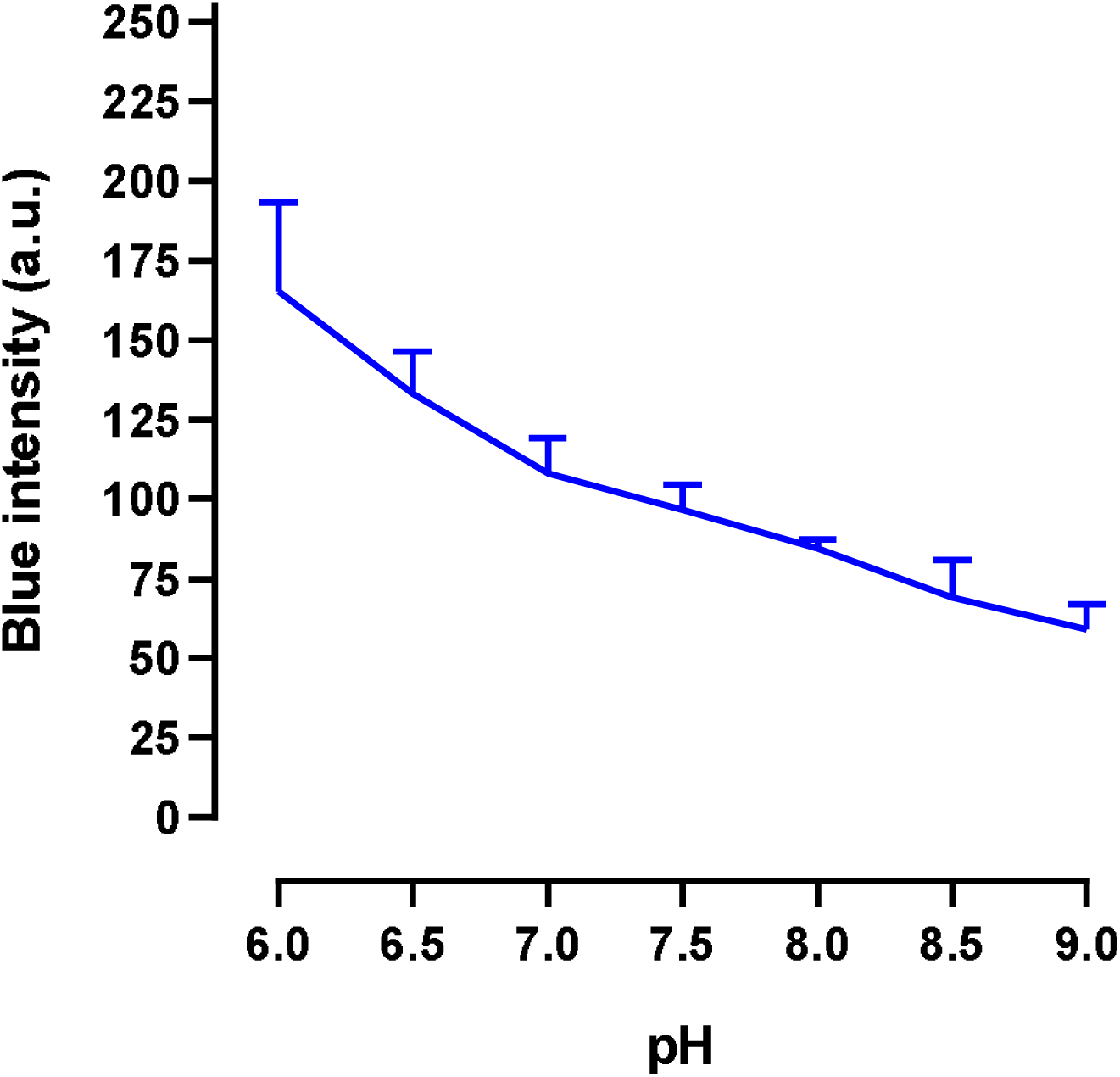
Arduino-recorded RGB absorbance of HPTS-loaded MPs at different pH values. Microparticles were incubated in MES isotonic buffer (50 mM) for pH 6.0-6.5 and TBS isotonic buffer (50 mM) for pH 7.0-9.0 for approx. 5 minutes before measurement. Results presented as mean ± SD (n=3).

#### 2.2.3. Detection of pH-sensitive hydrogel using Arduino-coupled RGB detector

To investigate if the pH-dependent color signal of the dye-loaded MPs can be detected in a hydrogel, we encapsulated them in a calcium-crosslinked alginate hydrogel and incubated them at pH 6.0 to 9.0. As in 2.1.3., the hydrogel’s color signals were detected *in vitro* in 6-well plates and *ex vivo* on full-thickness excisional dorsal mouse wounds. A decrease in blue signal intensity can be observed as pH values increased, both *in vitro* (**Figure 7**); the similarity in the decreases indicates that there is low interference by the background.

**Figure 7.**
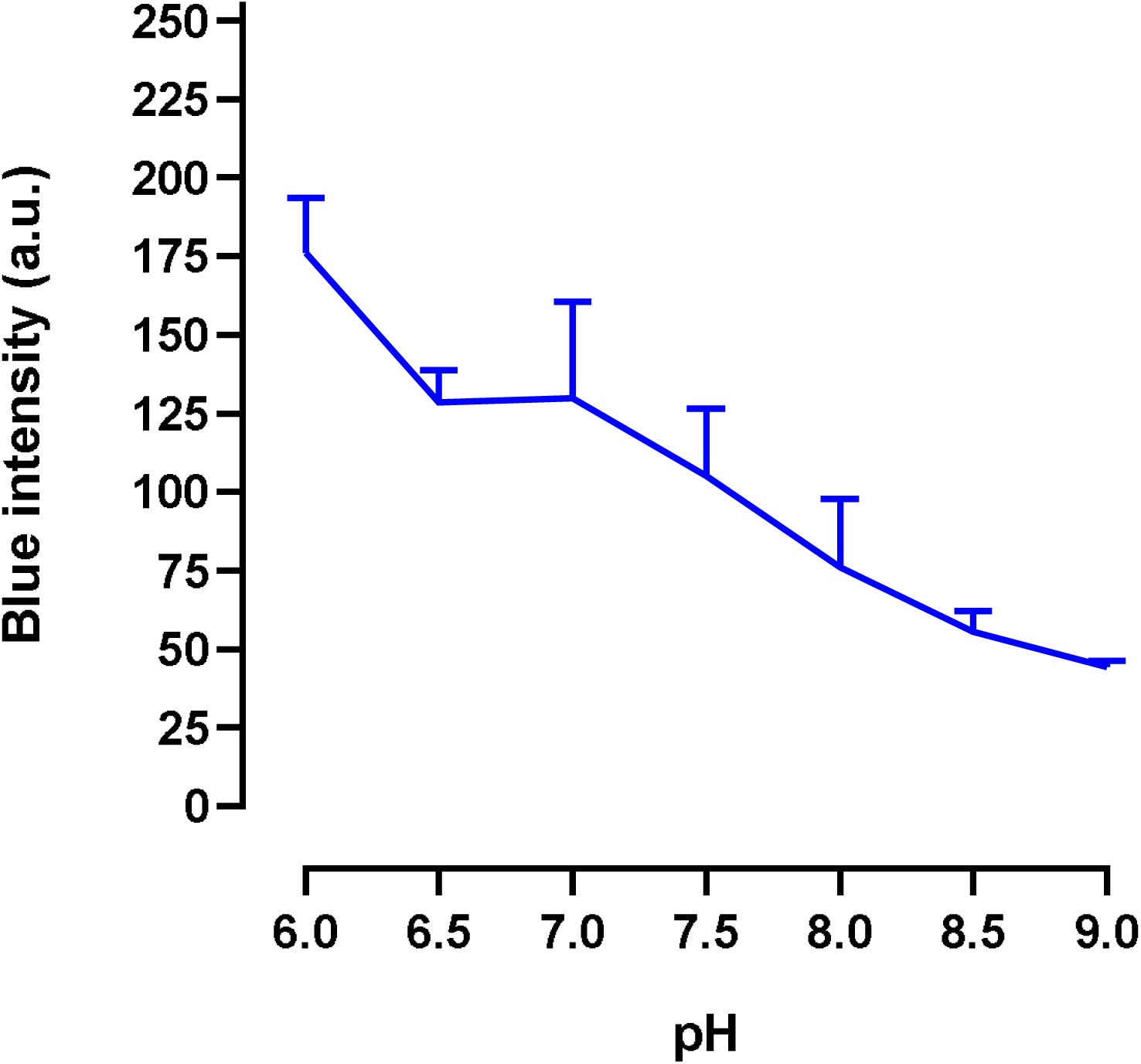
Arduino-recorded RGB absorbance of pyranine-loaded BTA microparticle-encapsulating calcium alginate hydrogels in vitro. Hydrogels were incubated in corresponding buffer solutions (MES isotonic buffer 50 mM for pH 6.0-6.5 and TBS isotonic buffer 50 mM for pH 7.0-9.0) for approx. 5 minutes before measurement. Results presented as mean ± SD (n=3).

## 3. Conclusion

DFUs are a serious and prevalent complication of diabetes. Current clinical guidelines recommend non-specific macroscopic evaluation of the wounds despite a growing understanding of diabetic wound pathophysiology. There is a need for new molecular diagnostics for improved disease staging, treatment selection, and assessment of treatment response. Here, we are presenting a colorimetric pH-sensitive bandage for wound diagnostics at the point-of-care, and validated two detection methods: a smartphone camera and a homebuilt Arduino-coupled RGB sensor. The pH sensing mechanism is based on a pH-sensitive dye that is loaded onto microparticles and physically immobilized in a calcium alginate hydrogel to minimize dye release. The hydrogel exhibited a colorimetric response to pH differences in the clinically relevant range *in vitro* and *ex vivo* on full thickness dorsal mouse wounds. We demonstrated that the colorimetric signal is visible and can be quantified rapidly using a commercial app of a smartphone camera and an Arduino-coupled RGB sensor. These findings encourage further development of this pH-sensing system for molecular diagnostics in DFUs.

## 4. Experimental Section

### Set-up of ColorMeter-based RGB sensor

The ColorMeter-based sensor relies on the camera of a smartphone (iPhone 13) with its integrated flash LED. Upon installation of the ColorMeter RGB Colorimeter App, pictures were taken with the flash LED from the iPhone’s incorporated camera app at zoom 2X. For reproducibility of measurements, the smartphone was mounted on a selfie stick at 20 cm height from the camera lens to the table. For RGB absorbance quantification, pictures were imported in the ColorMeter RGB Colorimeter App and the in-app focus point size was adjusted to cover the entirety of the sample (smaller circle for wells containing HPTS solutions, bigger squares for whole hydrogels) to produce an accurate reading of the average of the color contained in the region of interest (ROI) (**Figure S12**).

### Set-up of Arduino-based RGB sensor

The TCS34725 RGB sensor includes an integrated LED for illuminating the surface being measured. Adjacent to the sensor is the RGB detector, which features a total of 12 photodiodes arranged in a 3×4 matrix covered with a color-specific filter sensitive to red, green and blue wavelengths (**Figure S7**). Upon installing the Arduino IDE software, the TCS34725 RGB sensor was soldered to an array of 8-pin BergStick Connectors to be mounted on an 830 Tie-Points Breadboard and connected to an Arduino UNO Rev3 controller board using four Dupont wires (**Figure S8**). The 3v3 pin of the sensor, which supplies voltage, was connected to the 3v3 pin on the controller board with a red wire. The GND pin of the sensor, providing ground current, was linked to the GND pin on the controller board via a black wire. The SCL pin of the sensor, which handles current input, was connected to pin A5 on the controller board using an orange wire. The SDA pin, which manages both current input to the sensor and output to the controller board, was connected to pin A4 on the controller board with a green wire (Figure S8, **Figure S9**). Once the connections are finalized, the code was implemented using the `jmparatte/jm_LiquidCrystal_I2C` library, which includes additional libraries such as `Adafruit_TCS34725` by Adafruit, `LiquidCrystal` by Arduino, and `Wire.h`. These libraries are essential for managing the data input and output functions of the detector (**Figure S10**). When the sensor is directed at a sample, the photodiodes detect the reflected light and convert the light intensity into an electrical current, which is then processed by an ADC converter that transforms this current into voltage that the Arduino UNO Rev3 reads. The controller board processes these RGB readings according to the programmed code. The raw data intensities for individual R, G and B values are displayed in the Serial Monitor screen of the Arduino IDE software at 100-second intervals (**Figure S11**).

### Investigation of HPTS solutions for RGB detection

To investigate if the absorbance of the dye can be detected with a smartphone camera and with an Arduino-based RGB sensor, we prepared HPTS solutions at different concentrations and different pH values. To investigate pyranine detection at different concentrations, HPTS was reconstituted in PBS isotonic buffer (50 mM, pH 7.4) at 0, 10, 50, 100, 250 and 500 µM. The HPTS solutions were then transferred to a 96-well plate (BRANDplates®, 781965, flat white bottom) for RGB colorimetry. For smartphone camera analysis, pictures were taken with the default iPhone 13 Camera App; pictures were then imported in the ColorMeter RGB Colorimeter App and measurements were taken as the average color of each well, as described previously (Figure S12). For Arduino-based analysis, the TCS34725 sensor, fixed on the breadboard, was aligned directly on top of the wells with the help of the incorporated LED flash and measurements were taken as an average color of each well. To investigate pyranine detection at different pH values, HPTS was reconstituted at a concentration of 100 µM in MES isotonic buffer (50 mM) for pH values 6.0 and 6.5 and TBS isotonic buffer (50 mM) for pH values 7.0, 7.5, 8.0, 8.5 and 9.0; pH was adjusted using HCl 1 N or NaOH 1 N as needed. The HPTS solutions were then transferred to a 96-well plate plate (BRANDplates®, 781965, flat white bottom) for RGB colorimetry with both smartphone camera and Arduino-based RGB sensor, as described for the detection of HPTS at different concentrations.

### pH-sensitivity of HPTS-loaded BTA MPs for RGB detection

To investigate if pyranine loaded onto microparticles can be detected with a smartphone camera and with an Arduino-based RGB sensor, we analysed dye-loaded MPs at different concentrations and different pH values. To investigate HPTS-loaded MPs with the smartphone camera system, 150 mg of BTA MPs were incubated in 5 mL of pyranine at 0, 100 or 500 µM reconstituted in corresponding pH buffer solutions (MES for pH 6.0 and 6.5 and TBS for pH 7.0 to 9.0) for 15 minutes at 37 °C with agitation and protected from light. The dispersion was then purified from unbound dye by centrifugation (three times at 15 000 *x* g for 1 min). After purification, the MPs were put in corresponding buffers solutions again for 5 to 10 minutes in a 24-well plate (Sarstedt, 83.3921, flat bottom, transparent) for color signal stabilization and pictures were taken as described previously with the iPhone Camera app and imported on the ColorMeter App for RGB colorimetry. To investigate HPTS-loaded MPs with the Arduino-based RGB sensor, 150 mg of BTA MPs were incubated in 5 mL of pyranine at 100 µM in corresponding pH buffer solutions (MES for pH 6.0 and 6.5 and TBS for pH 7.0 to 9.0) for 15 minutes at 37 °C with agitation and protected from light. This concentration of HPTS (100 µM) was selected. After the loading, the dispersion was then purified from unbound dye by centrifugation (three times at 15 000 x g for 1 min). After purification, the MPs were put in corresponding buffers solutions for 5 to 10 minutes in a 24-well plate (Sarstedt, 83.3921, flat bottom, transparent) and RGB analysis was done by directing the TCS34725 sensor at each well, as described previously.

### pH-sensitivity of HPTS-loaded BTA MPs loaded into hydrogels

To investigate if the pH-dependent color signal of the dye-loaded MPs can be detected in a hydrogel, we first investigated the distribution of our MPs into the initial hydrogel matrix. In order to detect colorimetric signal all over the bandage, as we are measuring the average of color all over the sample as described previously, we encapsulated 5, 20, 40, 60, 100 and 150 mg of BTA MPs in our previously described ionically crosslinked calcium alginate hydrogels ^[8]^. The MPs were incubated in 5 mL of pyranine at 100 µM in PBS isotonic buffer (50mM, pH 7.4) for 15 minutes at 37 °C with agitation and protected from light for dye-loading. After purification of unbound dye by centrifugation (three times at 15 000 x g for 1 min), the MPs were encapsulated in 120 µL of medium viscosity sodium alginate 8 % in water and 40 µL of calcium sulfate dihydrate suspension 192 mM in water, as per our previous protocol ^[8]^. A quantity of 150 mg of BTA MPs was selected for one hydrogel since it had the most homogeneous distribution throughout the matrix (Figure S3), however, the alginate matrix was insufficient in total volume to fully contain all MPs. Following this, we doubled all other quantities in our formulation to better encapsulate the MPs by mixing them with 240 µL of medium viscosity sodium alginate 8 % in water and 80 µL of calcium sulfate dihydrate suspension 174 mM in water with final concentrations of 6% alginate and 43.5 mM calcium sulfate. After mixing, each hydrogel is shaped into disks with spatulas for post-crosslinking in a beaker of calcium chloride solution 25 mM (100 mL) for 4 hours at 37 °C. Following their preparation, pH sensitivity of the hydrogels was tested *in vitro* and *ex vivo* on both the smartphone camera system and Arduino-based system. For *in vitro* analysis, hydrogels were placed in pH-corresponding buffer solutions for 5-10 minutes for color stabilization in 6-well plates (Sarstedt, 83.3920, flat bottom, transparent) and RGB colorimetry was analyzed as described previously. For *ex vivo* analysis, excisional full thickness mouse wounds (C57BL/6 male mice, 18 weeks of age and Podocin-Adams17 female mice, 11 weeks of age) were performed using a biopsy punch with a diameter of 10 mm. Hydrogels were incubated in pH-corresponding buffer solutions for 5-10 minutes for color stabilization in 6-well plates. For each mouse, one hydrogel cut in 2 pieces was placed on the wounds (Figure S5) and RGB colorimetry analysis was performed as previously described.

### Statistical analysis

GraphPad was used for the statistical analysis.

## Supporting Information

Supporting Information is included at the end of this manuscript.

## Acknowledgements

SM gratefully acknowledges funding from Natural Sciences and Engineering Research Council of Canada (Discovery Grant RGPIN-2022-04384), Canada Foundation for Innovation (Fonds des leaders John-R.-Evans 42712), Fonds de recherche du Québec – Nature et technologies (Relève professorale 330153).

## Data Availability Statement

The data that support the findings of this study have been included as part of this article and the Supplementary Information.

## Supplementary Information

**Figure S1.**
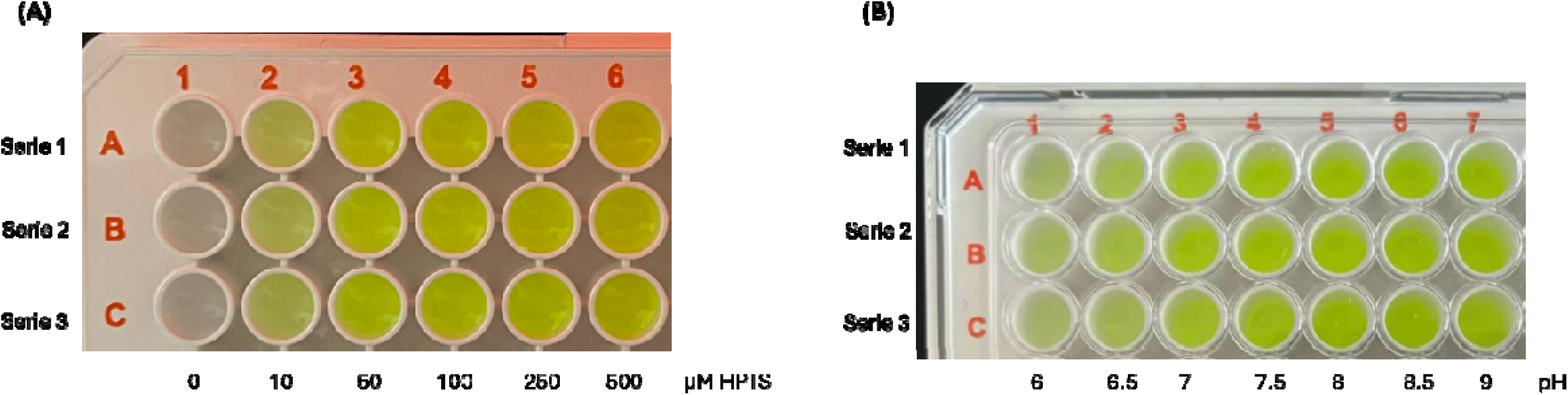
Pyranine (HPTS) solutions at varying concentrations and pH. **(A)** HPTS solutions at different concentrations (0-500 µM) in PBS isotonic buffer (50 mM, pH 7.4). **(B)** HPTS solutions at different pH values (6.0-9.0) in MES isotonic buffer (50 mM) for pH 6.0-6.5 and TBS isotonic buffer (50 mM) for pH 7.0-9.0.

**Figure S2.**
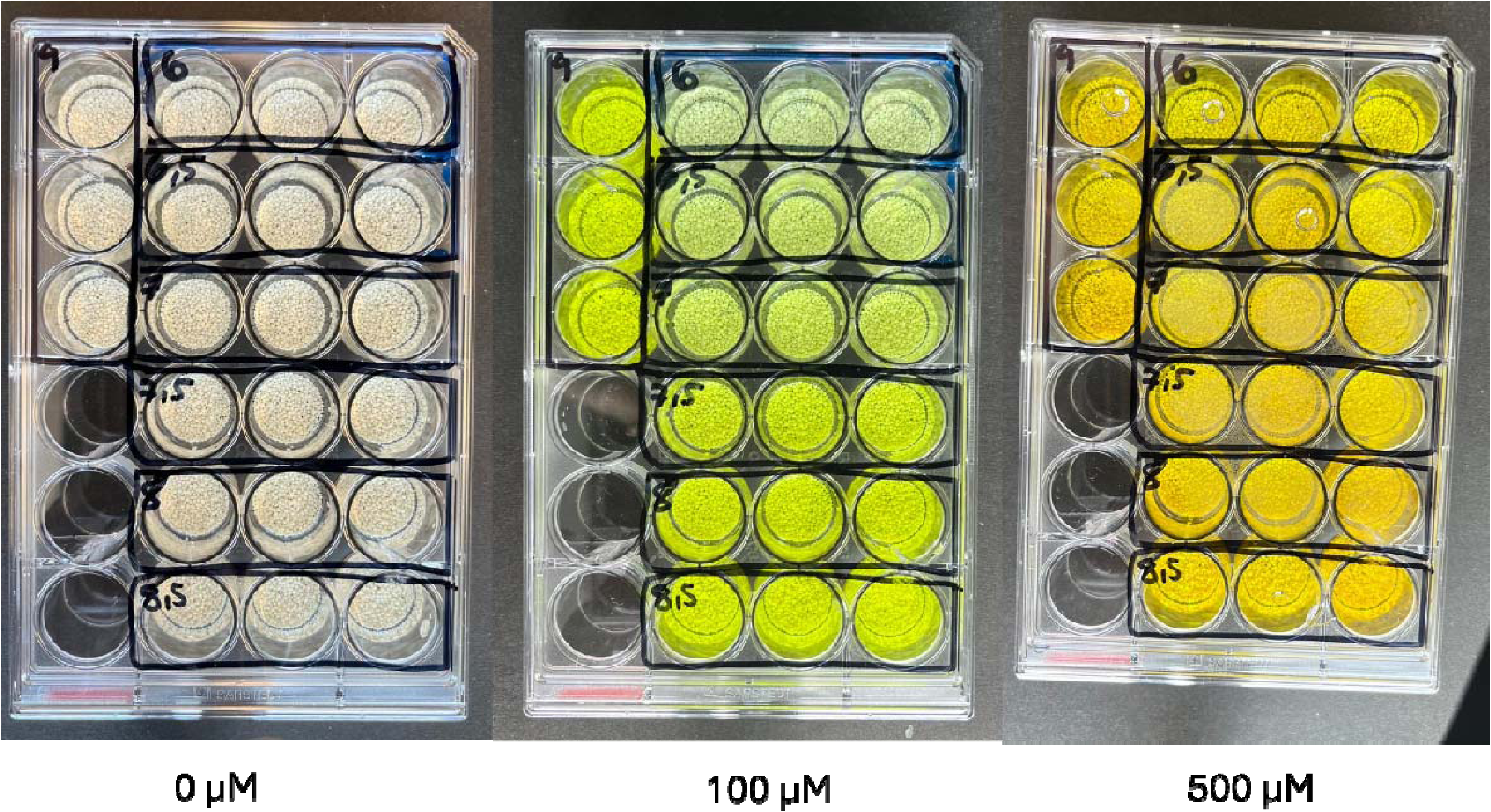
Microparticles loaded with pyranine (HPTS) at different concentrations and pH values. Microparticles were loaded with 0, 100, or 500 µM of HPTS and incubated in buffers ranging from pH 6.0 to 9.0 (MES isotonic buffer 50 mM for pH 6.0-6.5, TBS isotonic buffer 50 mM for pH 7.0-9.0).

**Figure S3.**
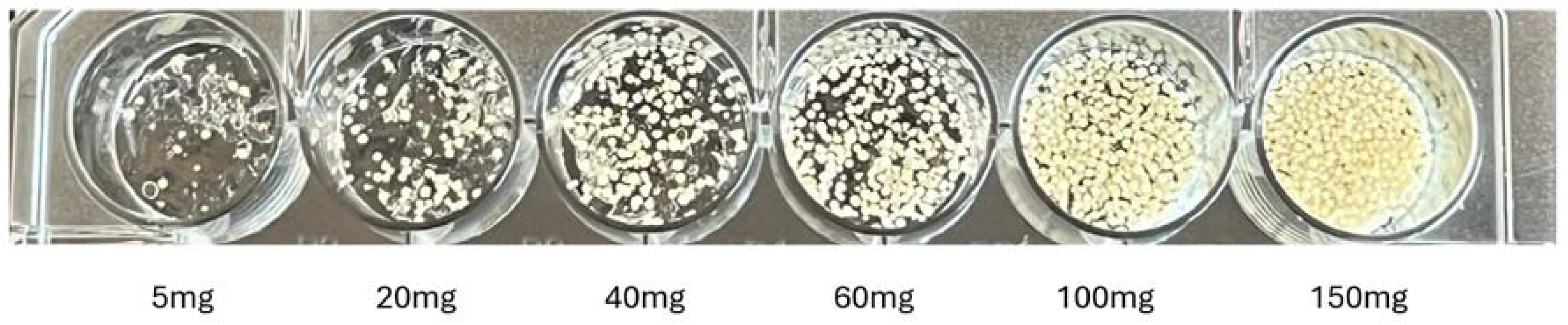
Hydrogels loaded with varying quantities of microparticles. Hydrogels were prepared with different amounts of microparticles to optimize their distribution in the matrix.

**Figure S4.**
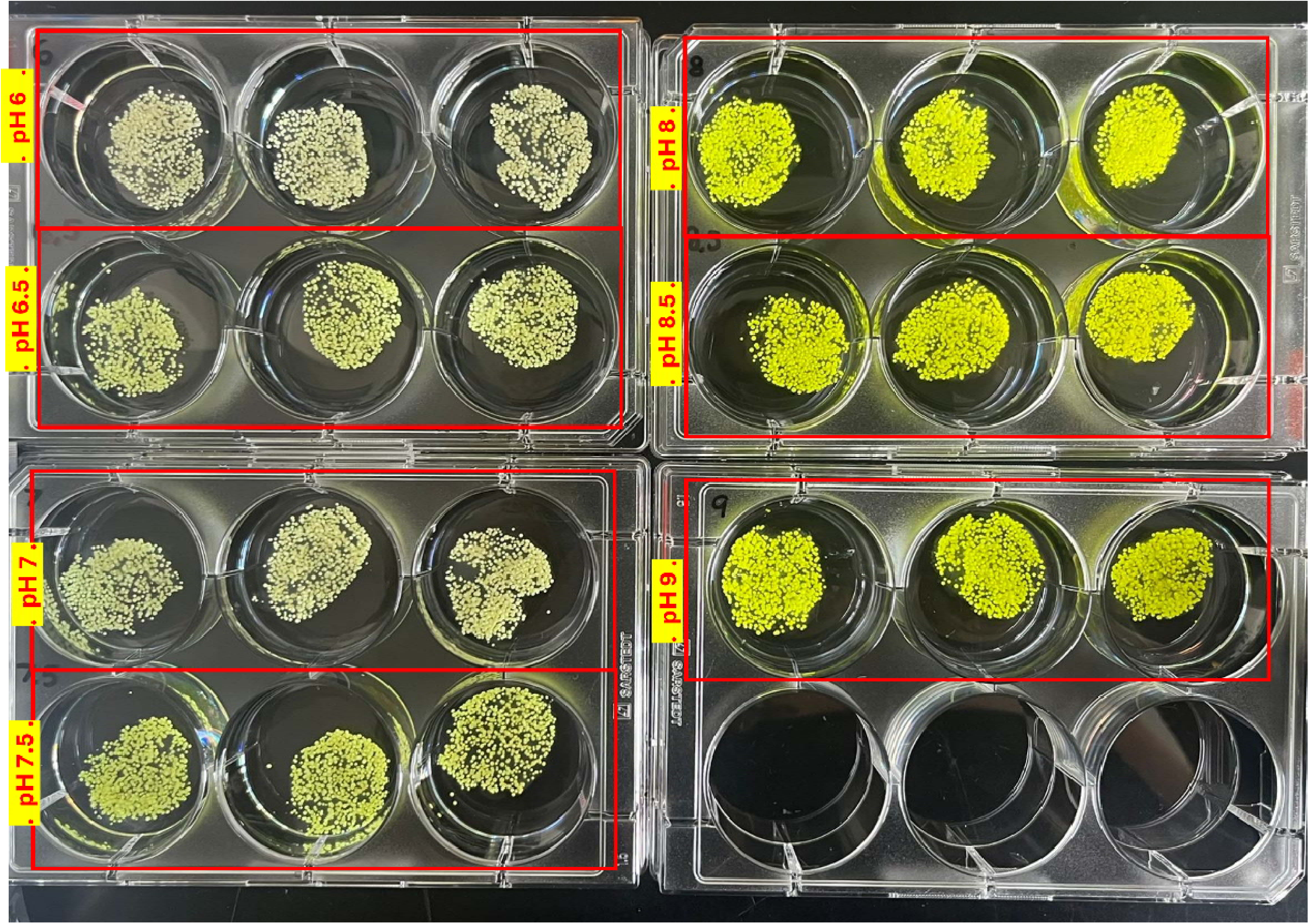
pH-responsive hydrogels at different pH values. Hydrogels were incubated in MES isotonic buffer (50 mM) for pH 6.0-6.5 and TBS isotonic buffer (50 mM) for pH 7.0-9.0 to demonstrate pH-dependent color changes.

**Figure S5.**
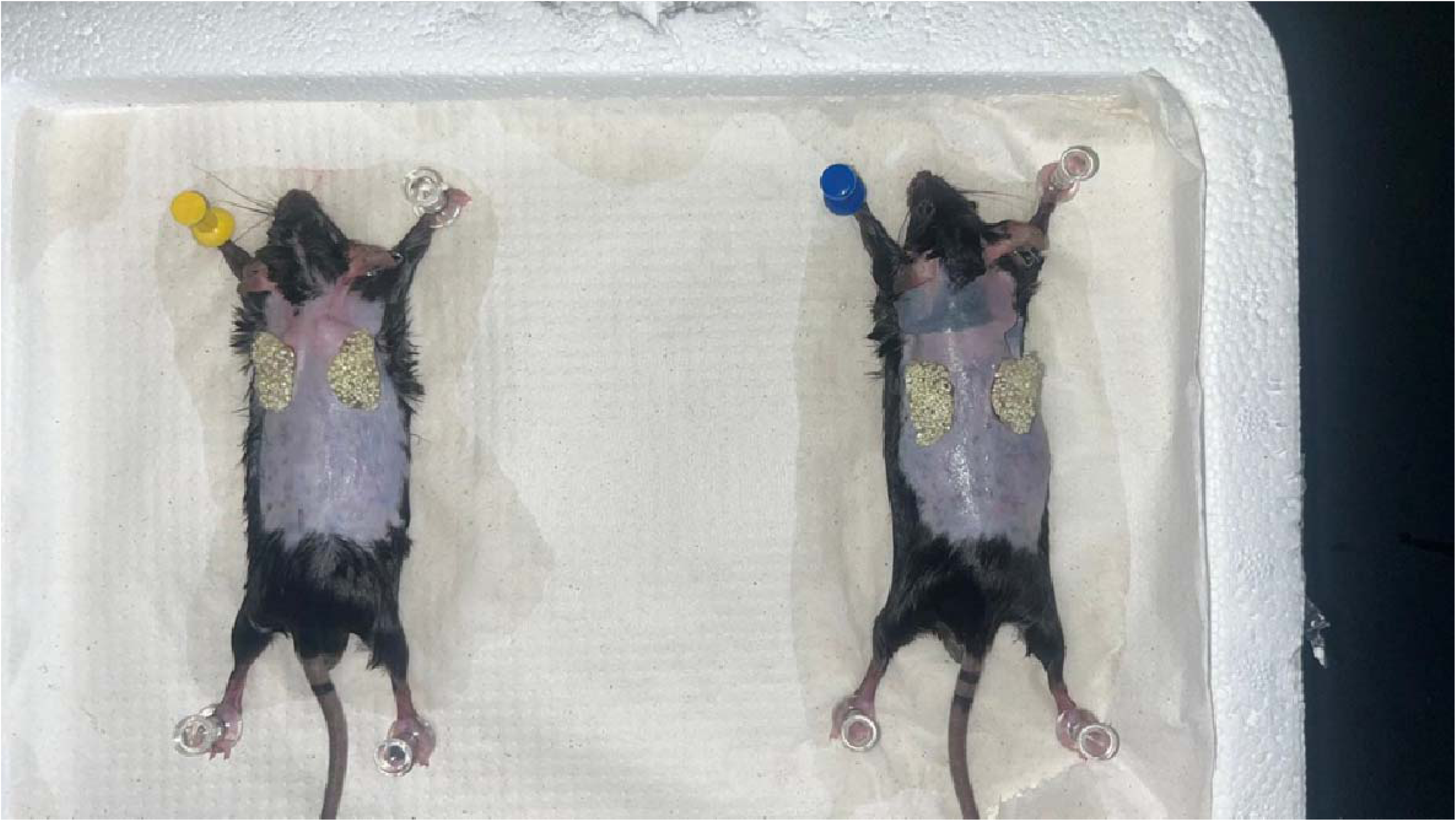
*Ex vivo* full-thickness excisional dorsal mouse wound model (2 out of 3 shown) with pH-sensitive hydrogels at pH 6.0.

**Figure S6.**
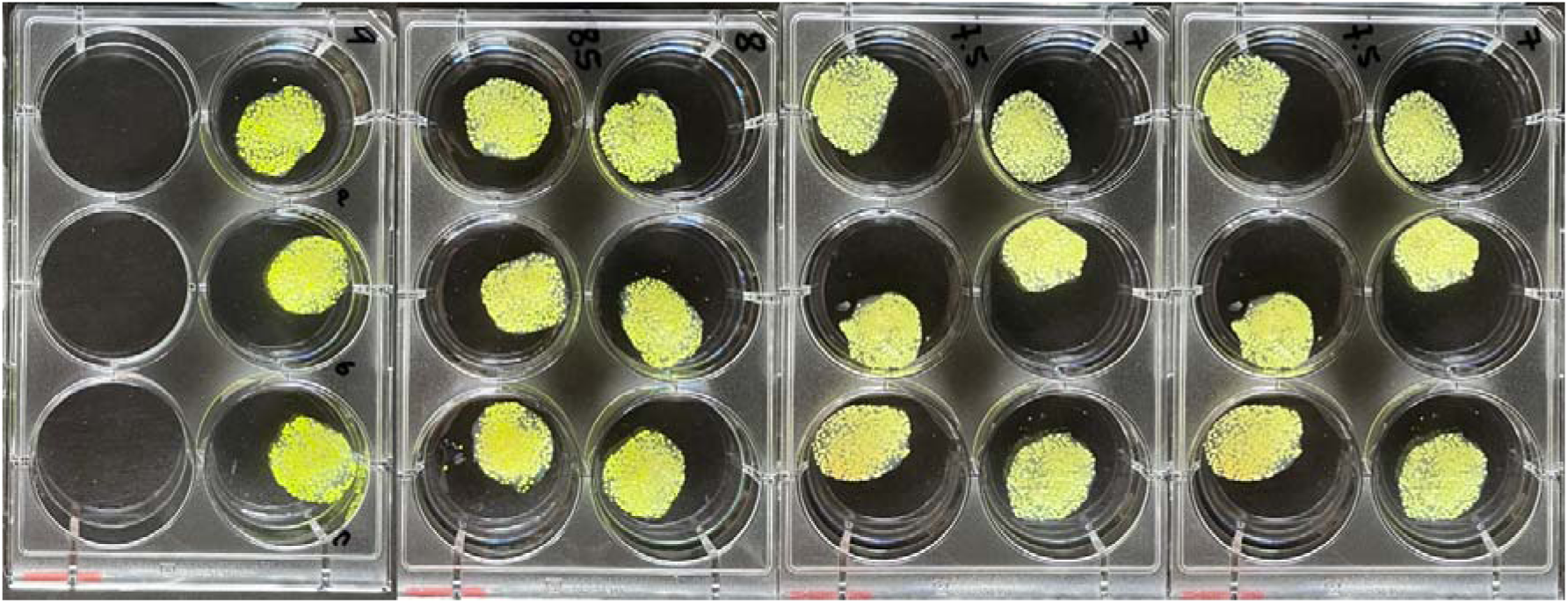
Short-term kinetics of pH-sensing hydrogels in vitro. Color changes of hydrogels were monitored at pH 6-9 over a 1-2 minute period to assess rapid response characteristics.

**Figure S7.**
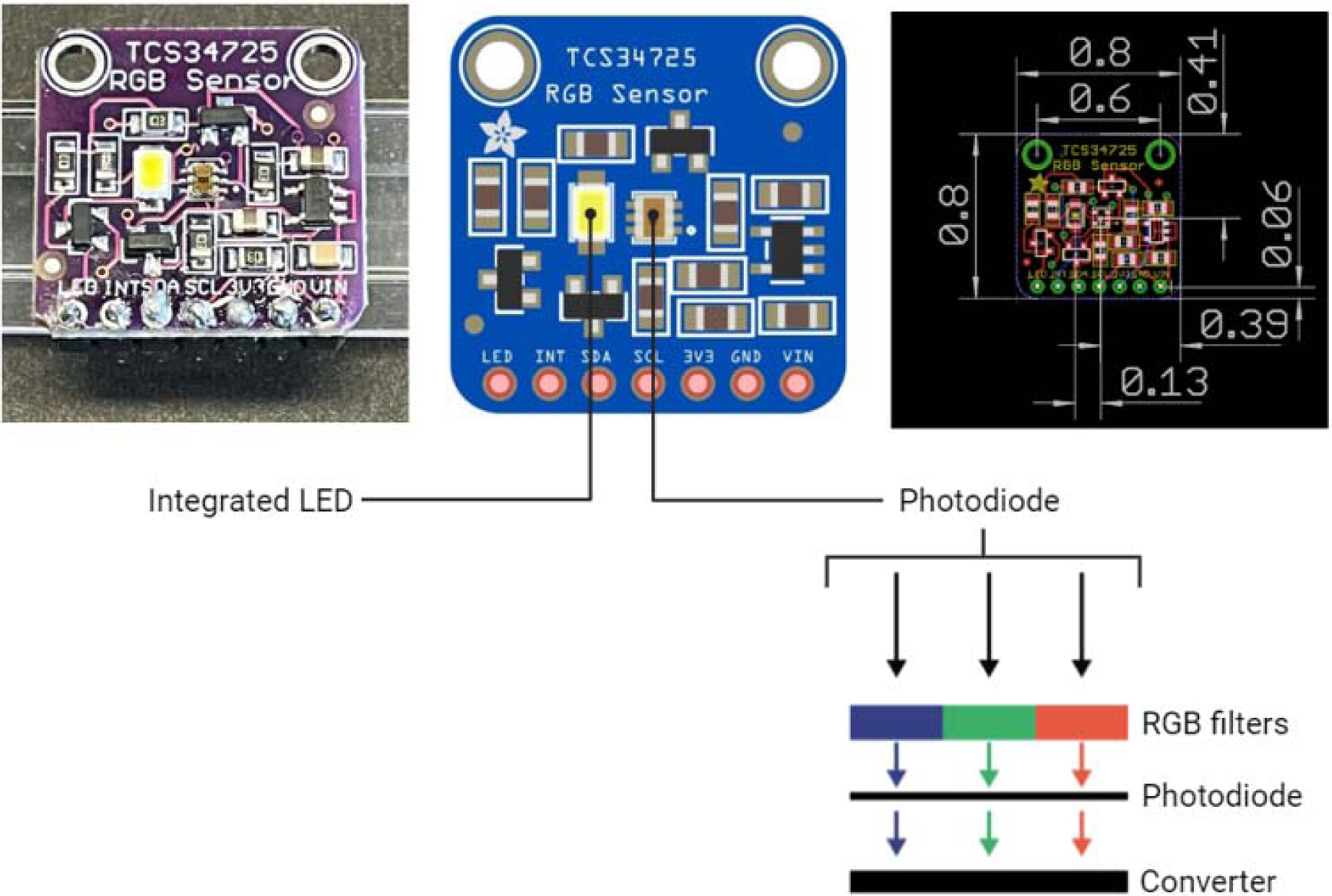
Technical specifications of the TCS34725 RGB sensor. Schematic diagram and dimensions (in millimeters) of the sensor used for colorimetric measurements.

**Figure S8.**
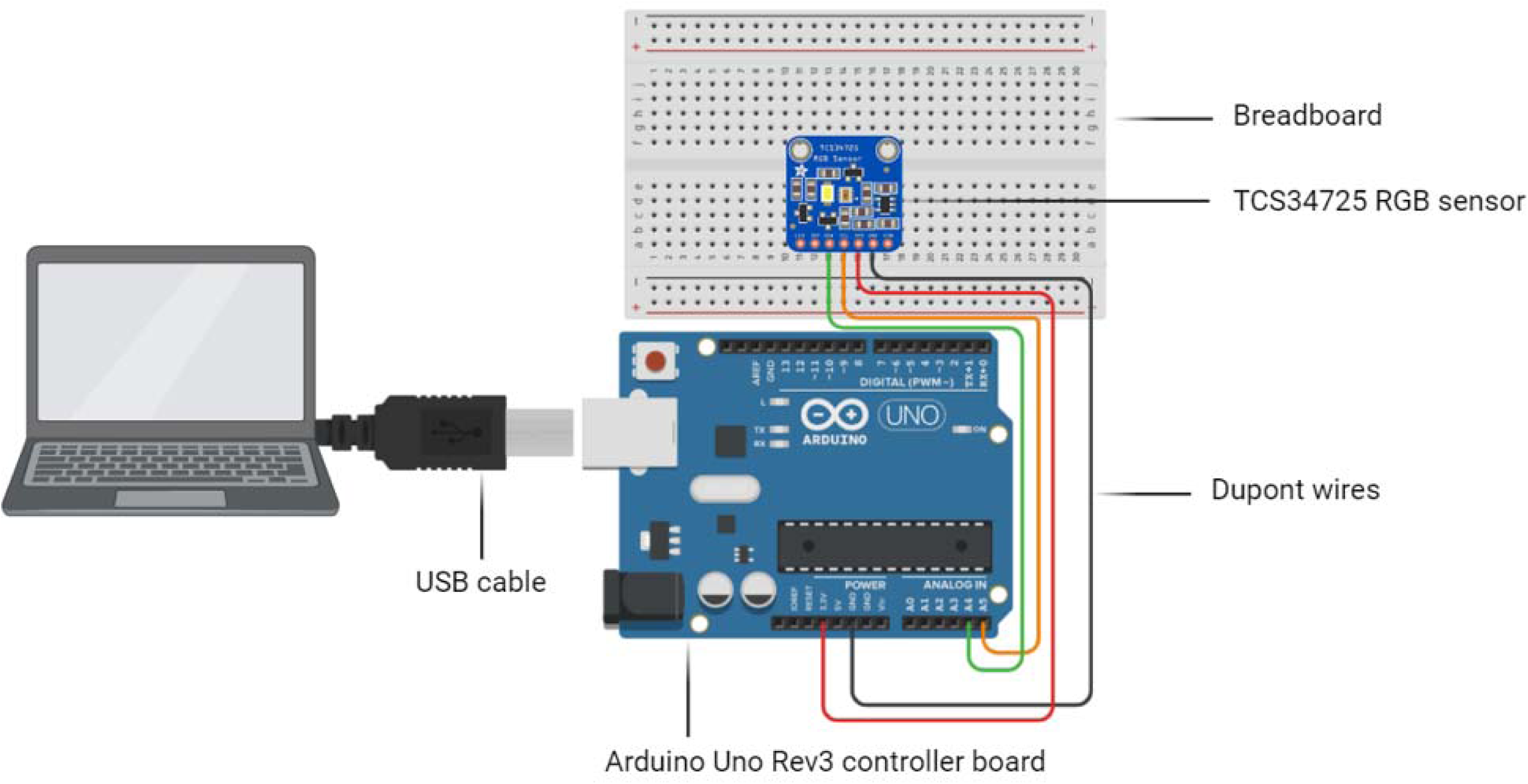
Connection diagram for TCS34725 RGB sensor and Arduino UNO Rev3. Schematic showing the breadboard connections between the RGB sensor and Arduino microcontroller.

**Figure S9.**
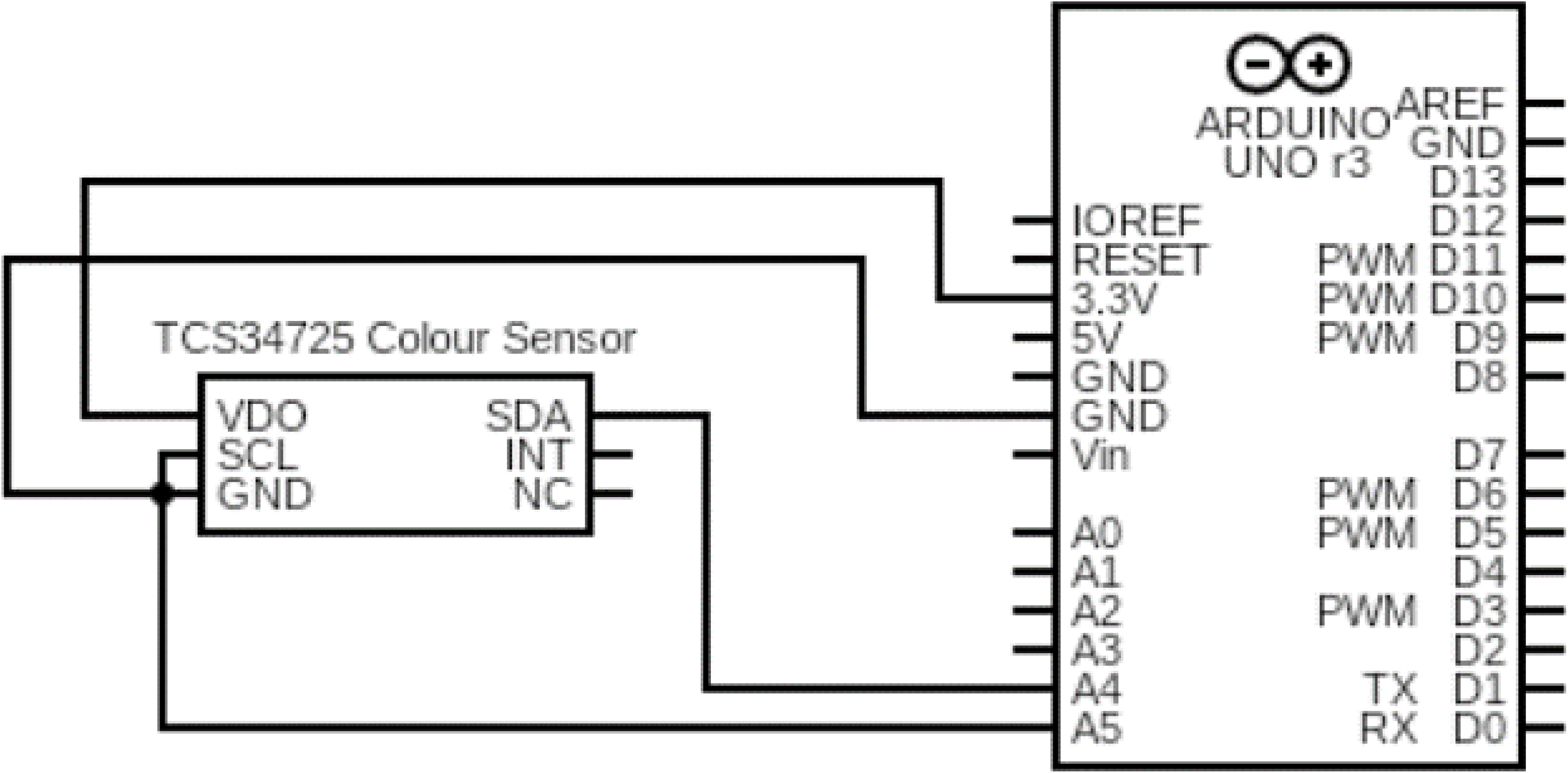
Circuit diagram of the Arduino RGB Detector. Detailed circuit schematic illustrating the connections between the TCS34725 RGB sensor and Arduino UNO Rev3 controller.

**Figure S10.**
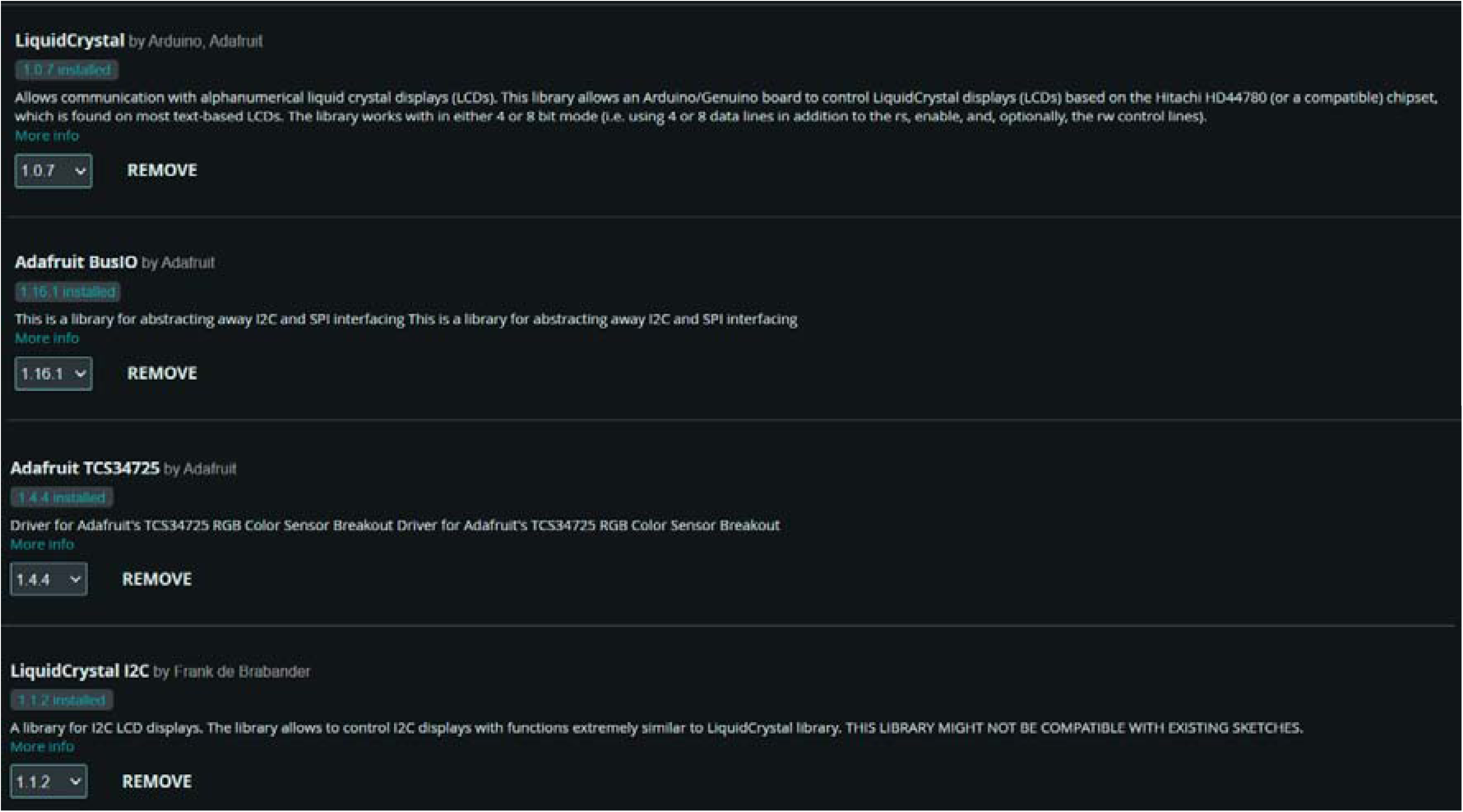
Arduino IDE 2.3.2 libraries. List of essential libraries downloaded and installed in the Arduino Integrated Development Environment for the RGB detection system.

**Figure S11.**
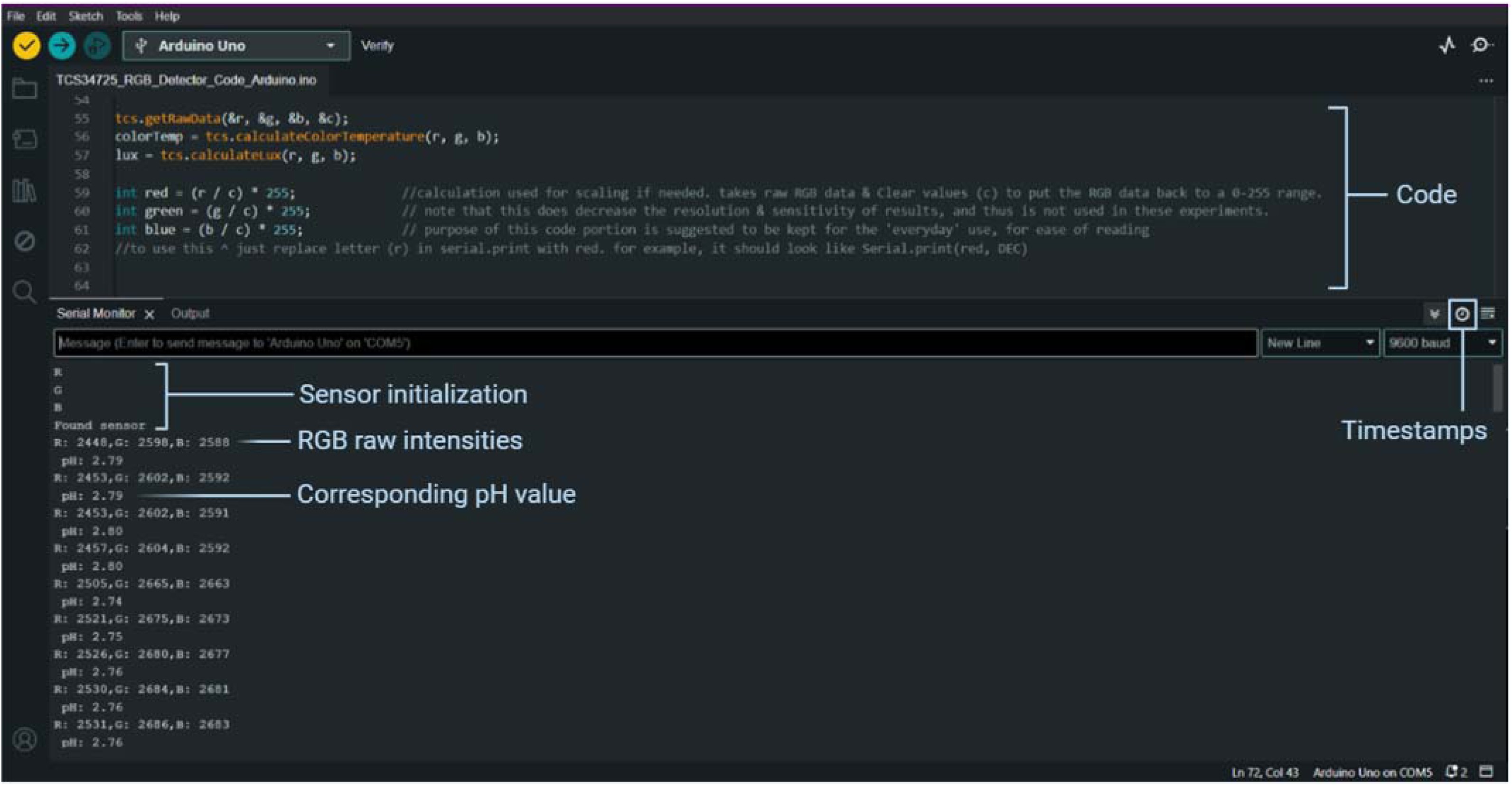
Arduino IDE software interface. Overview of the Arduino IDE during analysis, showing key features and code execution.

**Figure S12.**
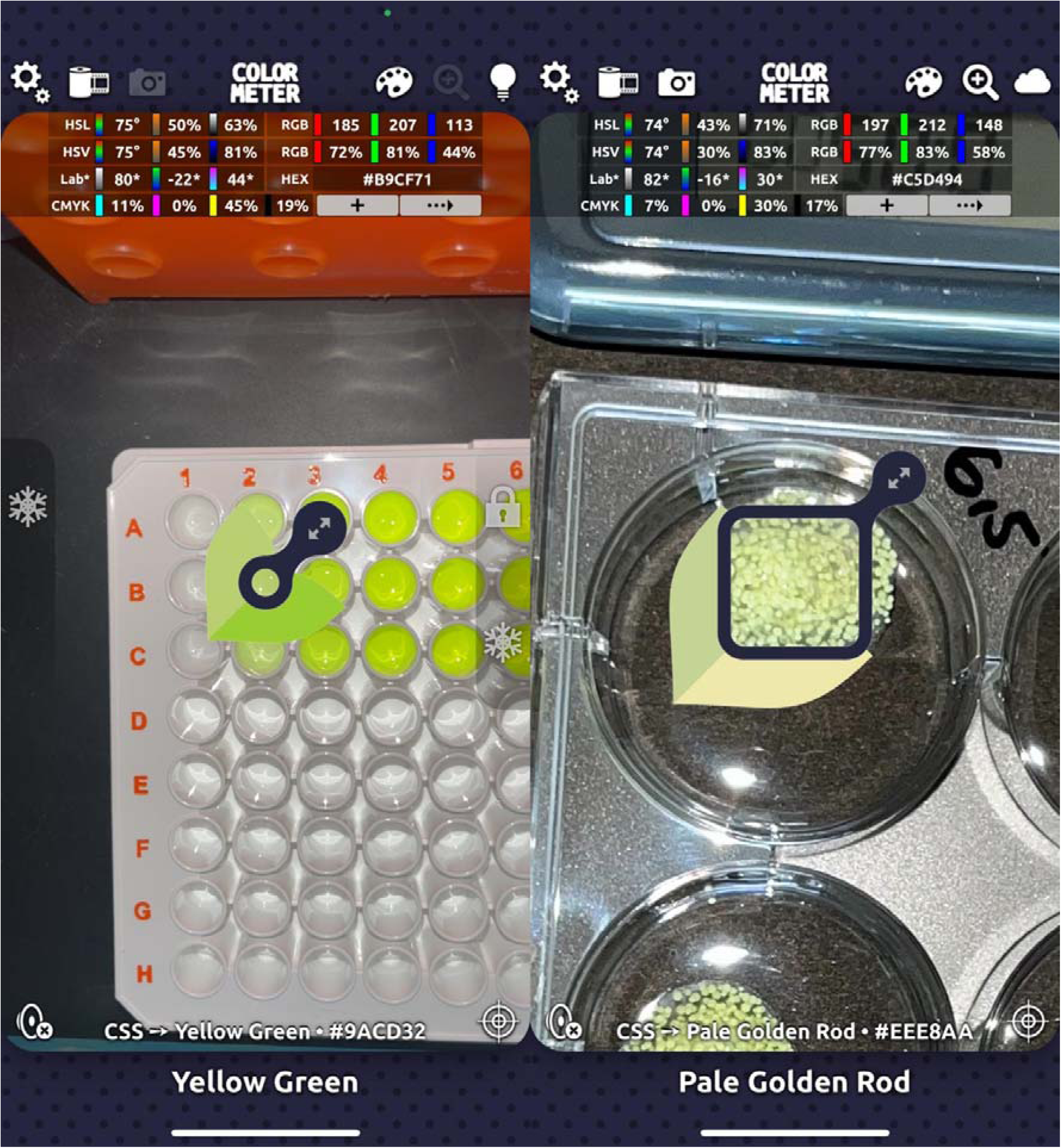
ColorMeter RGB Colorimeter App user interface on iPhone 13. **(Left)** RGB quantification of an HPTS solution sample. **(Right)** RGB quantification of a whole hydrogel. The focus point size can be adjusted to fit sample dimensions. Individual R, G, and B values are displayed in the top right corner of the screen. Images were captured using the iPhone’s camera app and imported into the ColorMeter App for stable measurements.

**Figure S13.**
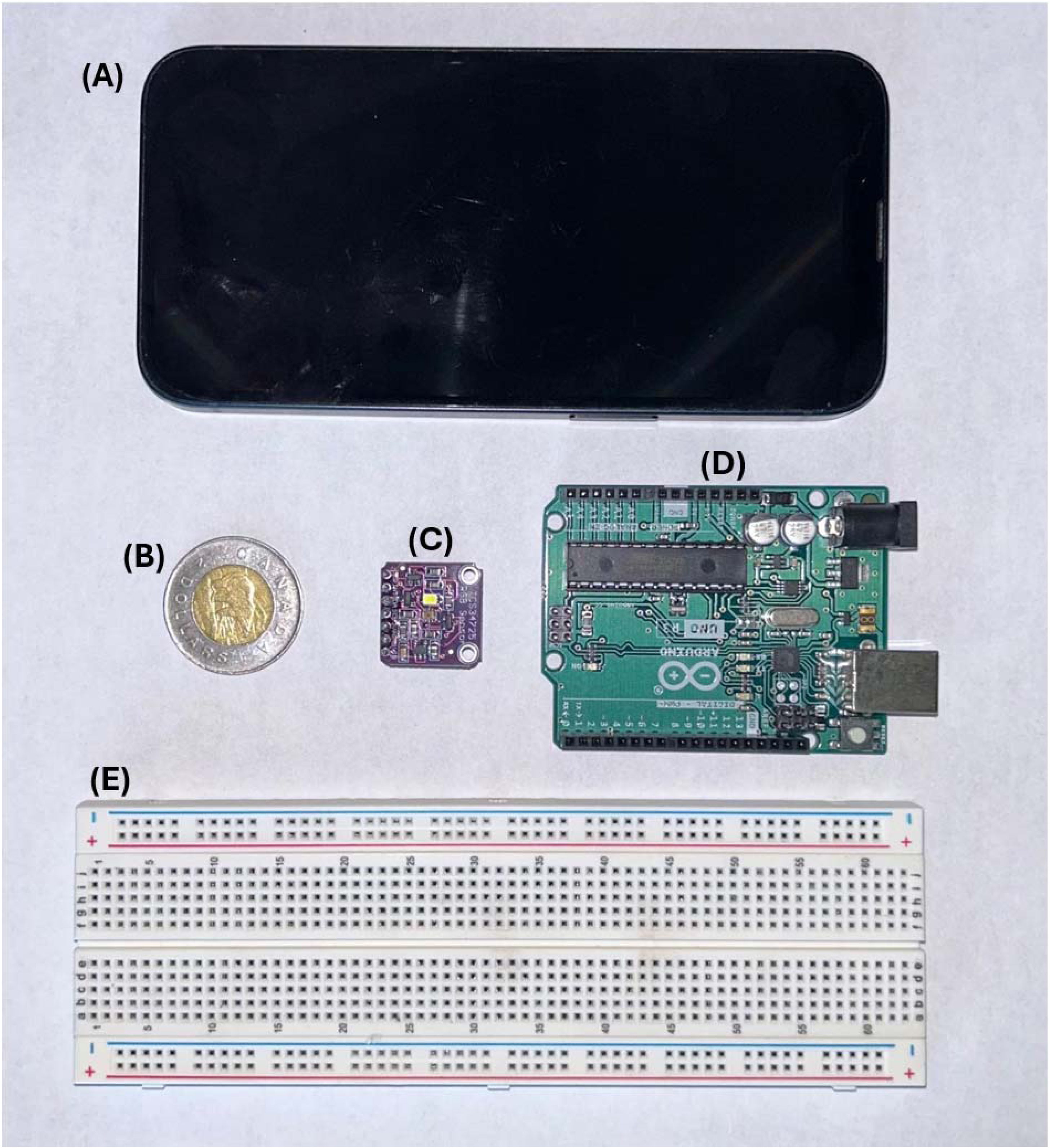
Comparison of pH sensing devices and components with a Canadian two-dollar coin (toonie). The image shows side-by-side comparisons of **(A)** an iPhone 13 smartphone used for ColorMeter app measurements, **(B)** a Canadian two-dollar coin (toonie) included for size reference, **(C)** the Arduino TCS34725 RGB color sensor chip, **(D)** the Arduino UNO Rev3 motherboard, and **(E)** the breadboard used for the Arduino-based pH detection system setup.

