## Supplementary Information for "Colorimetric detection methods of pH-sensing wound dressing for point-of-care wound diagnostics"

Université de Montréal

2940 Chemin de Polytechnique, Montreal, QC H3T 1J4

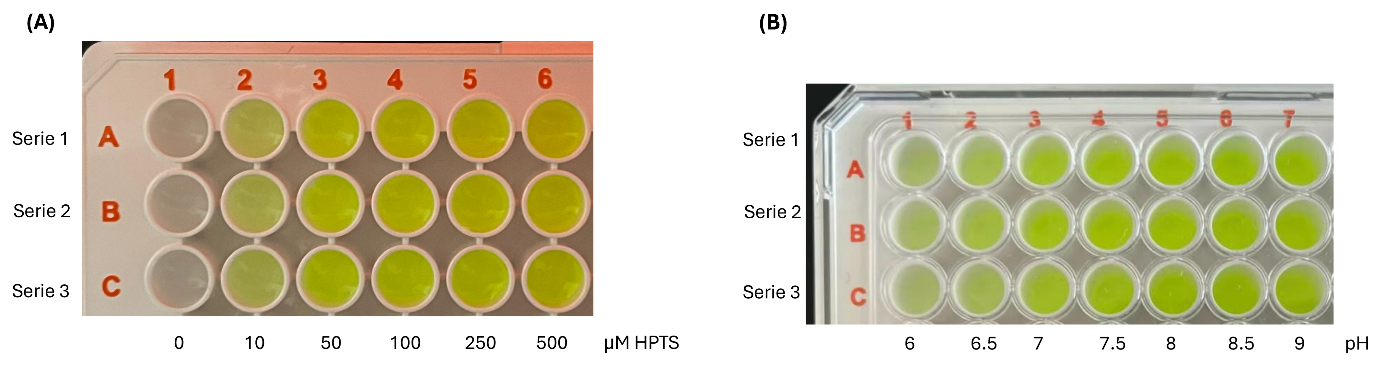

**Figure S1**. Pyranine (HPTS) solutions at varying concentrations and pH. **(A)** HPTS solutions at different concentrations (0-500 µM) in PBS isotonic buffer (50 mM, pH 7.4). **(B)** HPTS solutions at different pH values (6.0-9.0) in MES isotonic buffer (50 mM) for pH 6.0-6.5 and TBS isotonic buffer (50 mM) for pH 7.0-9.0.

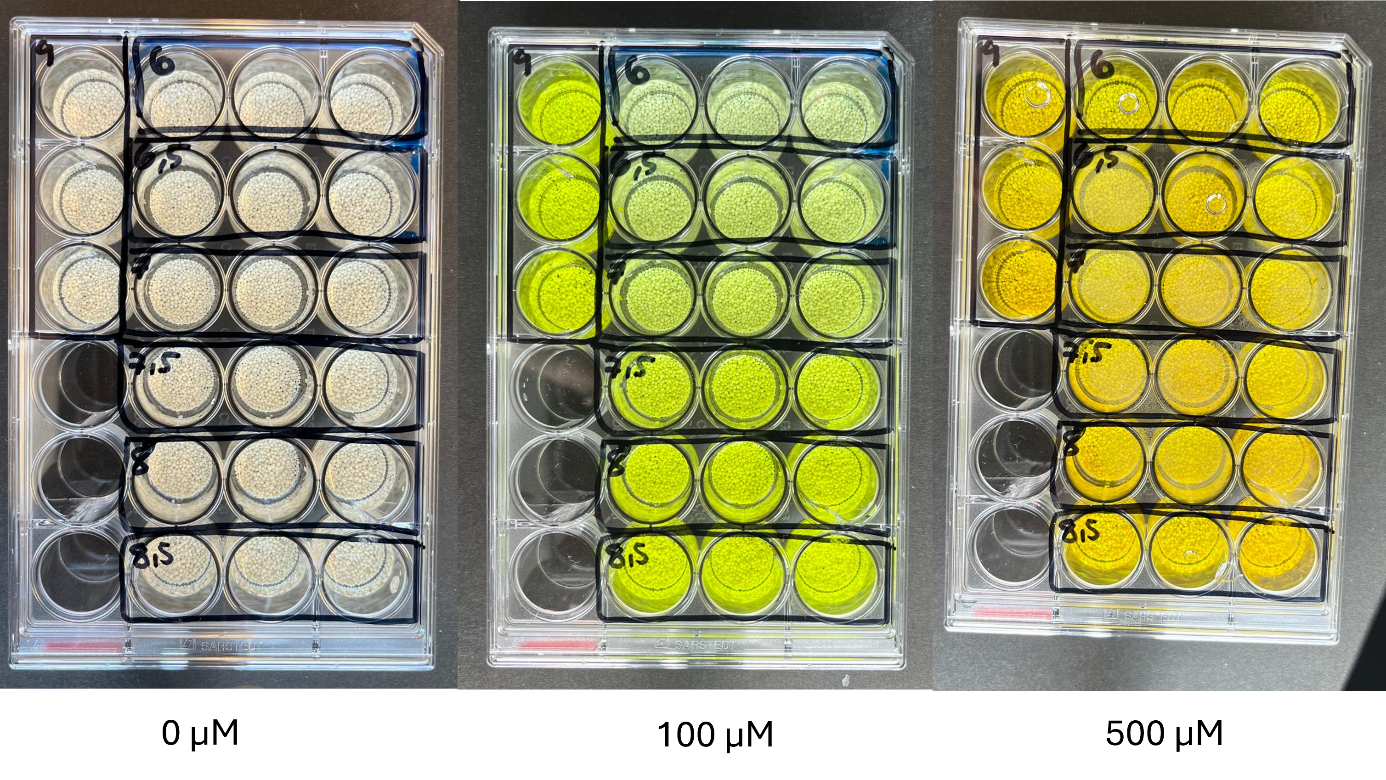

**Figure S2**. Microparticles loaded with pyranine (HPTS) at different concentrations and pH values. Microparticles were loaded with 0, 100, or 500 µM of HPTS and incubated in buffers ranging from pH 6.0 to 9.0 (MES isotonic buffer 50 mM for pH 6.0-6.5, TBS isotonic buffer 50 mM for pH 7.0-9.0).

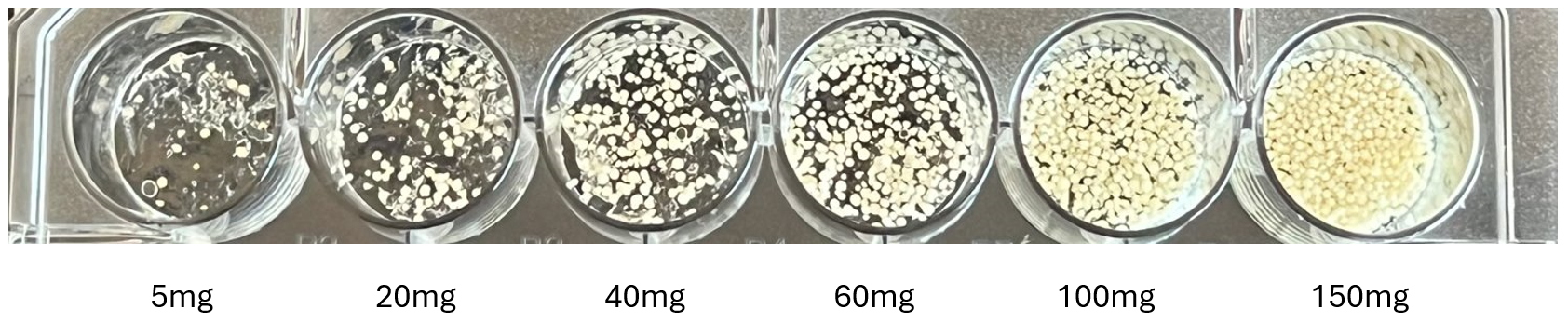

**Figure S3**. Hydrogels loaded with varying quantities of microparticles. Hydrogels were prepared with different amounts of microparticles to optimize their distribution in the matrix.

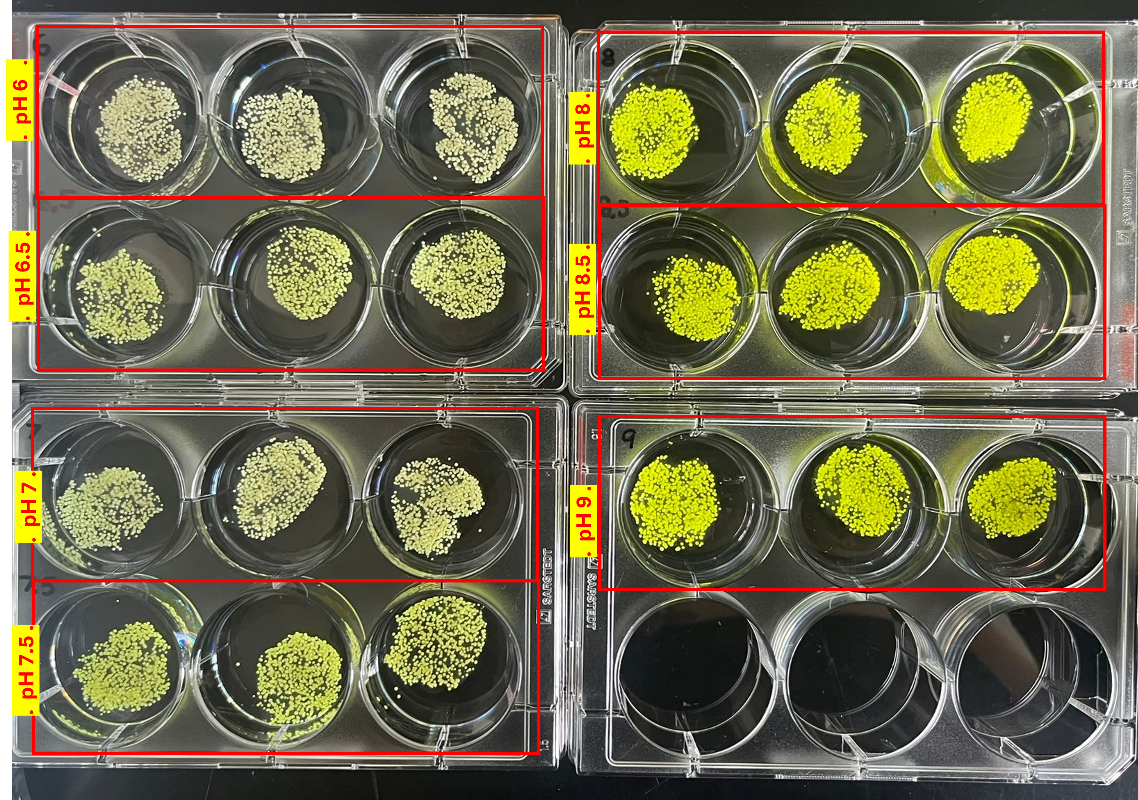

**Figure S4**. pH-responsive hydrogels at different pH values. Hydrogels were incubated in MES isotonic buffer (50 mM) for pH 6.0-6.5 and TBS isotonic buffer (50 mM) for pH 7.0-9.0 to demonstrate pH-dependent color changes.

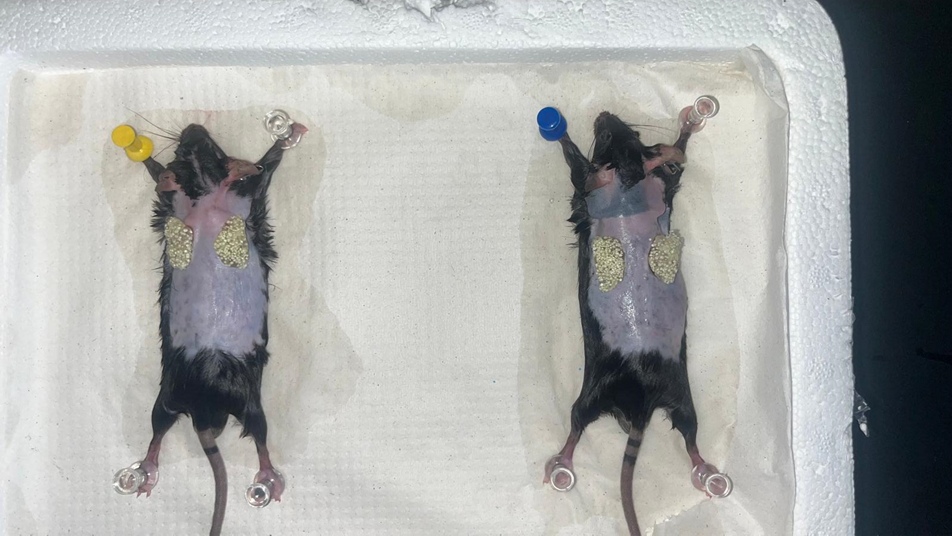

**Figure S5**. *Ex vivo* full-thickness excisional dorsal mouse wound model (2 out of 3 shown) with pH-sensitive hydrogels at pH 6.0.

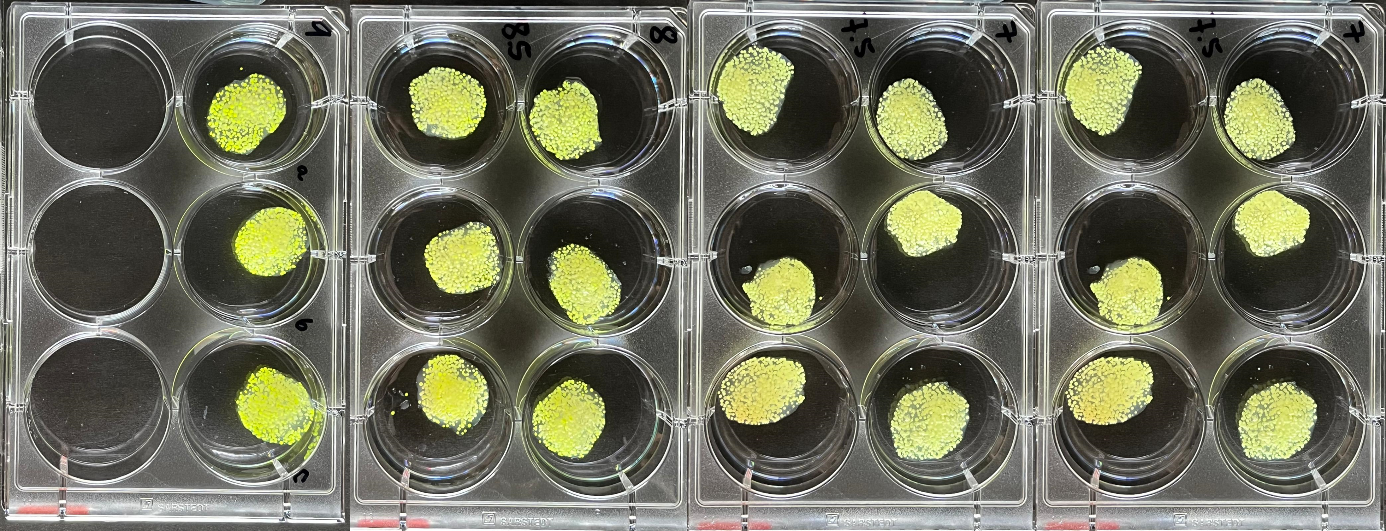

**Figure S6**. Short-term kinetics of pH-sensing hydrogels in vitro. Color changes of hydrogels were monitored at pH 6-9 over a 1-2 minute period to assess rapid response characteristics.

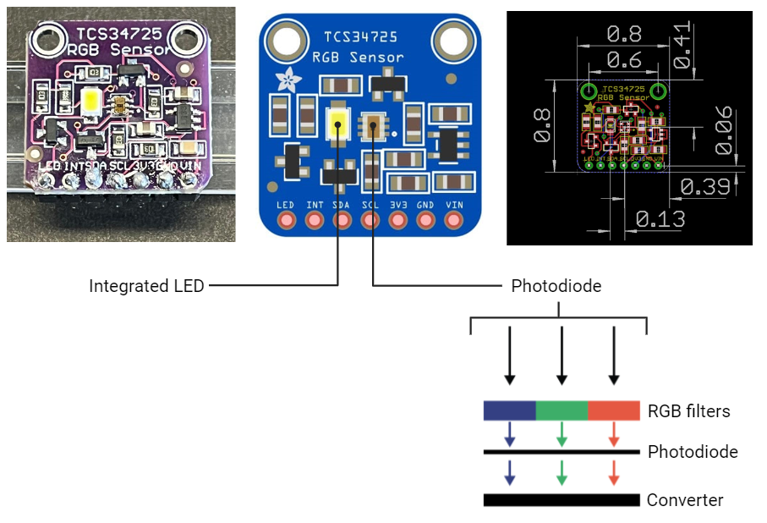

**Figure S7**. Technical specifications of the TCS34725 RGB sensor. Schematic diagram and dimensions (in millimeters) of the sensor used for colorimetric measurements.

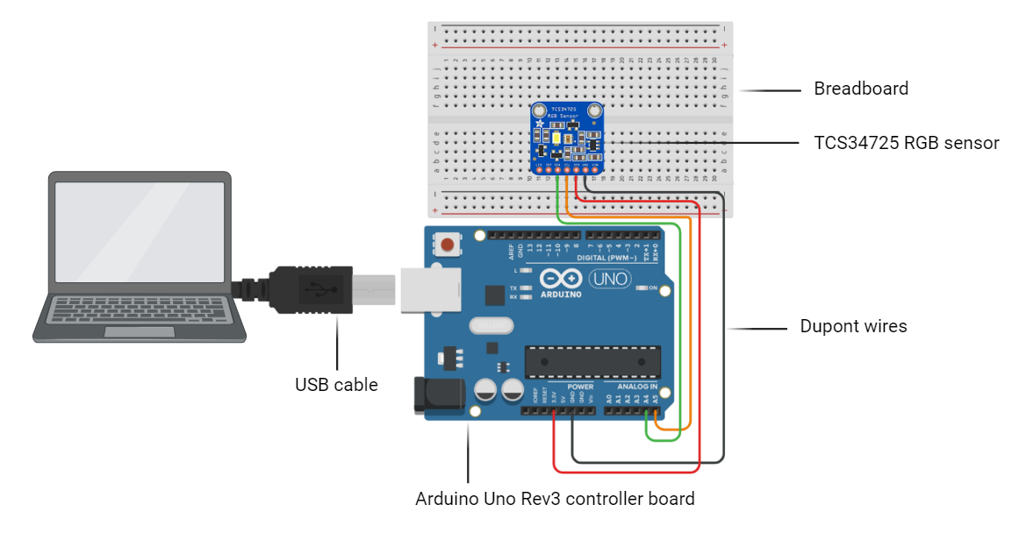

**Figure S8**. Connection diagram for TCS34725 RGB sensor and Arduino UNO Rev3. Schematic showing the breadboard connections between the RGB sensor and Arduino microcontroller.

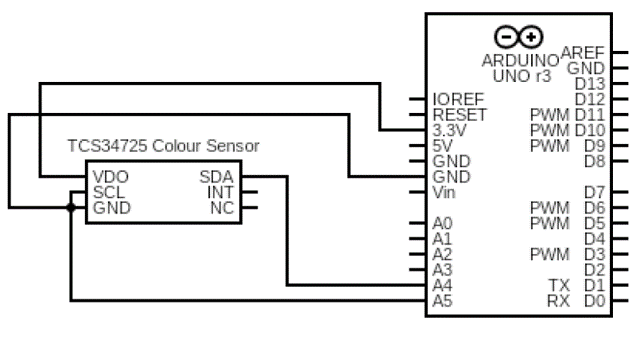

**Figure S9**. Circuit diagram of the Arduino RGB Detector. Detailed circuit schematic illustrating the connections between the TCS34725 RGB sensor and Arduino UNO Rev3 controller.

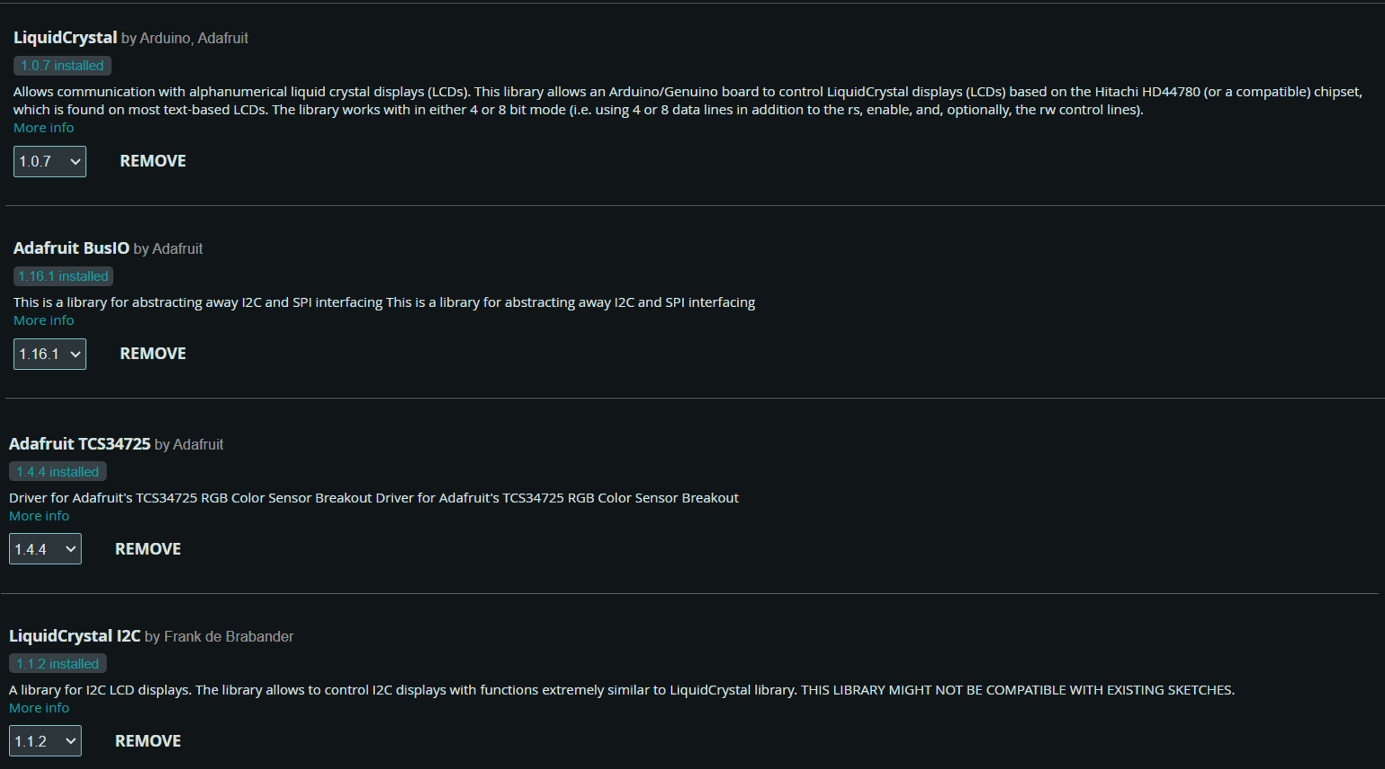

**Figure S10**. Arduino IDE 2.3.2 libraries. List of essential libraries downloaded and installed in the Arduino Integrated Development Environment for the RGB detection system.

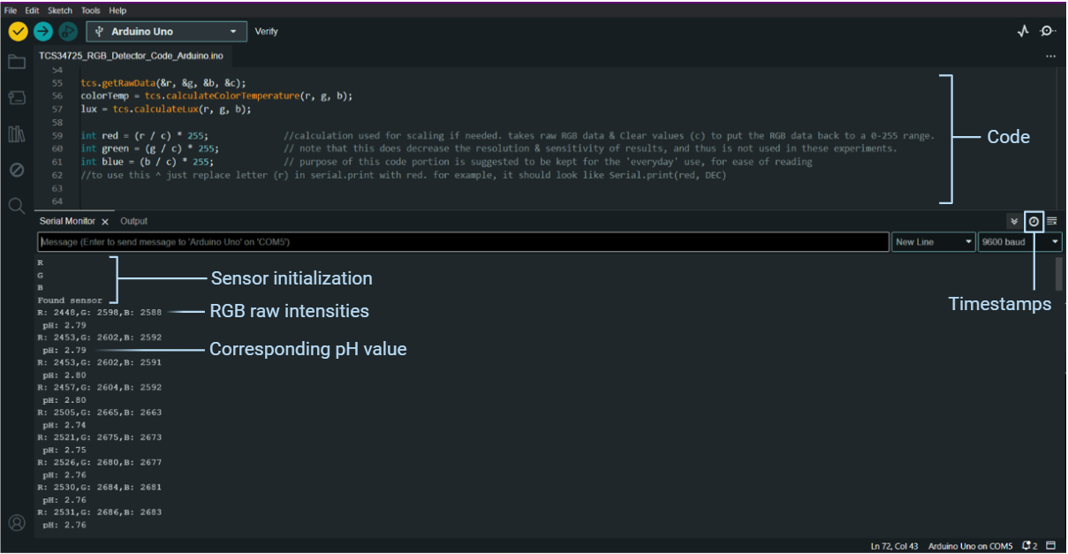

**Figure S11**. Arduino IDE software interface. Overview of the Arduino IDE during analysis, showing key features and code execution.

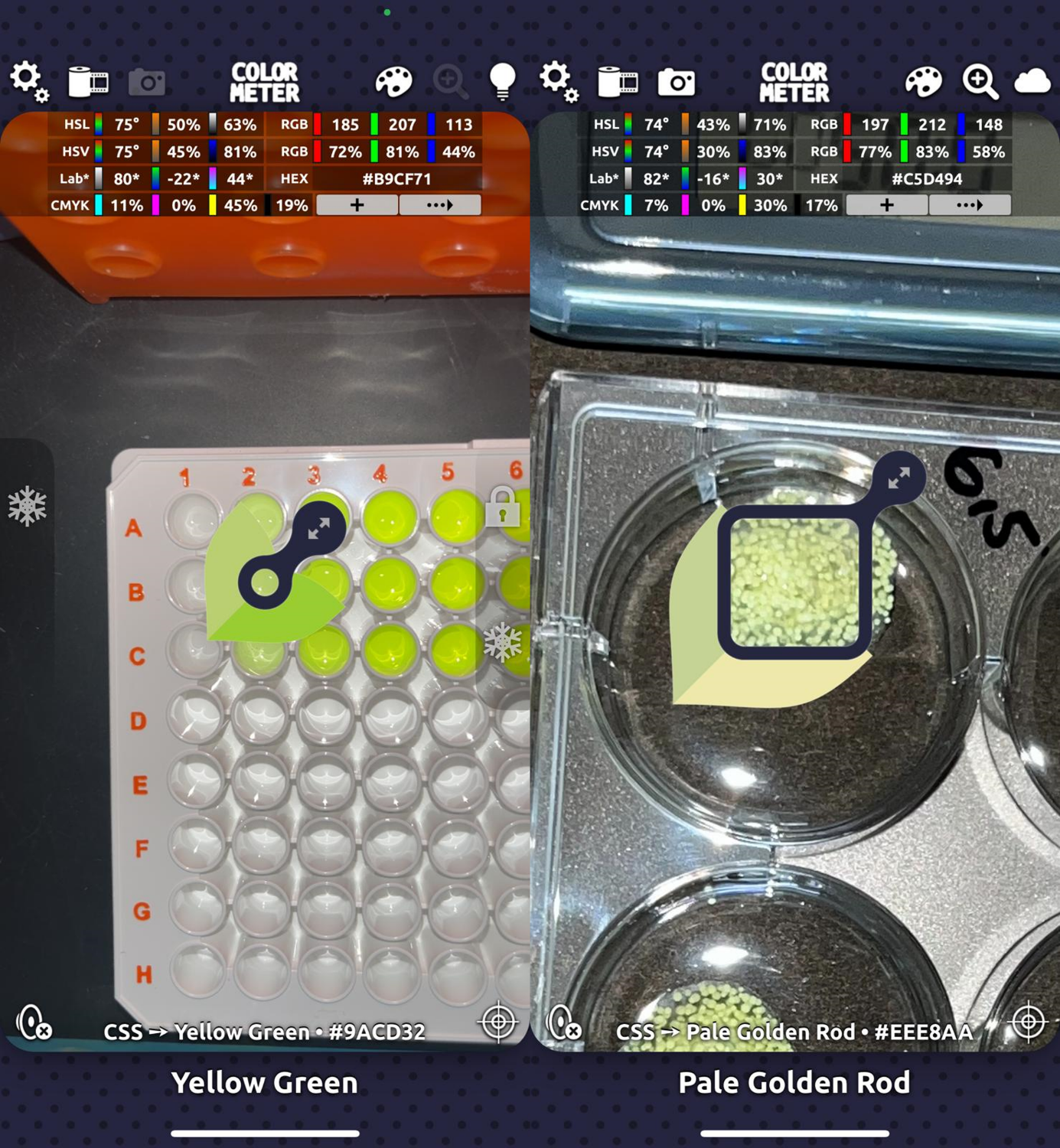
**Figure S12**. ColorMeter RGB Colorimeter App user interface on iPhone 13. **(Left)** RGB quantification of an HPTS solution sample. **(Right)** RGB quantification of a whole hydrogel. The focus point size can be adjusted to fit sample dimensions. Individual R, G, and B values are displayed in the top right corner of the screen. Images were captured using the iPhone's camera app and imported into the ColorMeter App for stable measurements.

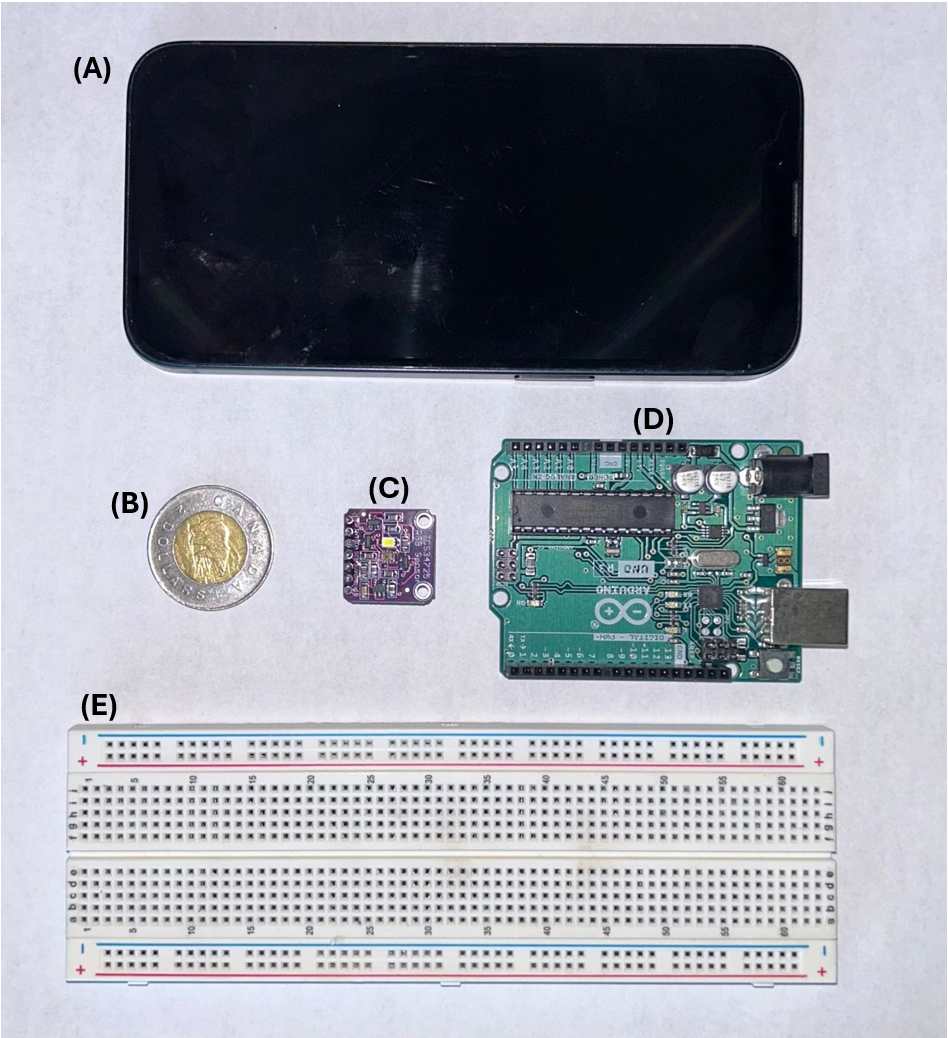

**Figure S13.** Comparison of pH sensing devices and components with a Canadian two-dollar coin (toonie). The image shows side-by-side comparisons of **(A)** an iPhone 13 smartphone used for ColorMeter app measurements, **(B)** a Canadian two-dollar coin (toonie) included for size reference, **(C)** the Arduino TCS34725 RGB color sensor chip, **(D)** the Arduino UNO Rev3 motherboard, and **(E)** the breadboard used for the Arduino-based pH detection system setup.
